# Multi-site phosphorylation of yeast Mif2/CENP-C promotes inner kinetochore assembly

**DOI:** 10.1101/2021.11.04.467375

**Authors:** Stephen M. Hinshaw, Yun Quan, Jiaxi Cai, Ann L. Zhou, Huilin Zhou

## Abstract

Kinetochores control eukaryotic chromosome segregation by connecting chromosomal centromeres to spindle microtubules. Duplication of centromeric DNA necessitates kinetochore disassembly and subsequent reassembly on the nascent sisters. To search for a regulatory mechanism that controls the earliest steps of kinetochore assembly, we studied Mif2/CENP-C, an essential basal component. We found that Polo-like kinase (Cdc5) and Dbf4-dependent kinase (DDK) phosphorylate the conserved PEST region of Mif2/CENP-C and that this phosphorylation directs inner kinetochore assembly. Mif2 phosphorylation promotes kinetochore assembly in a reconstituted biochemical system, and it strengthens Mif2 localization at centromeres in cells. Disrupting one or more phosphorylation sites in the Mif2-PEST region progressively impairs cellular fitness and sensitizes cells to microtubule poisons. The most severe Mif2-PEST mutations are lethal in cells lacking otherwise non-essential Ctf19 complex factors. These data suggest that multi-site phosphorylation of Mif2/CENP-C is a robust switch that controls inner kinetochore assembly, ensuring accurate chromosome segregation.

## Introduction

Passage through the cell cycle requires successive waves of kinase activity that coordinate transitions between the stages. This is particularly important at centromeres, which are the chromosomal sites of microtubule attachment. Kinetochores, the multiprotein assemblies that mediate this connection, are a nexus of kinase activity, acting both to supply and respond to the signals. In doing so, kinetochores negotiate cell cycle transitions by changing their shape and composition to accommodate DNA replication, tension sensing during metaphase, and chromosome movement during anaphase. While kinase activities required for regulated progression from metaphase through anaphase have been identified, those that enable kinetochore assembly during the earlier stages of the cell cycle have not.

The modular protein complexes that make up kinetochores can be designated as either inner or outer components (Biggins, 2013). Outer kinetochores mediate and respond to microtubule contact. Inner kinetochores connect the outer components to centromeric DNA and can be further subdivided into two groups: the Ctf19 complex (Ctf19c, or <u>C</u>onstitutive <u>C</u>entromere <u>A</u>ssociated <u>N</u>etwork (CCAN) in vertebrates) and a second complex containing the centromeric nucleosome with its essential adaptor protein, Mif2/CENP-C (hereafter Mif2) (Hara and Fukagawa, 2020; Hinshaw and Harrison, 2018; McKinley and Cheeseman, 2016; Musacchio and Desai, 2017). Of the 13 Ctf19c proteins, at least seven are phosphorylated *in vivo* (Okp1/CENP-Q, Ame1/CENP-U, Ctf19/CENP-P, Mcm21/CENP-O, Nkp1, Chl4/CENP-N, and Cnn1/CENP-T) (Bohm et al., 2021). Mif2 and Cse4/CENP-A, the histone H3-like component of the centromeric nucleosome, are also phosphorylated (Boeckmann et al., 2013; Westermann et al., 2003). Whether and how these post-translational modifications influence kinetochore assembly during the cell cycle are important open questions.

We and others have reported structures of inner kinetochore protein assemblies (Hinshaw and Harrison, 2019; 2020; Kixmoeller et al., 2020; Yan et al., 2019). With one notable exception (discussed below) (Ariyoshi et al., 2021), these structures show contacts between rigid components. They do not directly address post-translational protein modifications, which predominate on flexible N-terminal extensions not visible in the reported density maps. Concurrently, improved biochemical methods for kinetochore reconstitution from cell extracts have shown that cell cycle stage and, presumably, the associated kinase activities, influence robust inner kinetochore assembly (Lang et al., 2018). A mechanism for inner kinetochore stabilization by cell cycle-dependent kinase activity has not been defined and might vary from species to species. Compounding this uncertainty, multiple groups have presented divergent structural models for Ctf19c/CCAN-Cse4/CENP-A binding (Allu et al., 2019; Chittori et al., 2018; Hinshaw and Harrison, 2019; Pentakota et al., 2017; Yan et al., 2019; Zhou et al., 2021). Incomplete information about which inner kinetochore posttranslational modifications drive cell cycle-dependent reconfigurations, when the modifications occur, and whether they are conserved between species prevents contextualization of these proposals.

Mif2/CENP-C supports inner kinetochore assembly by linking Ctf19c/CCAN components to the centromeric nucleosome. It is essential for viability in both yeast and vertebrate cells (Brown et al., 1993; Kwon et al., 2007). Vertebrate CENP-C has two motifs that bind Cse4/CENP-A: a central domain and a C-terminal CENP-C motif (Carroll et al., 2010; Kato et al., 2013). The two sites are thought to be functionally redundant in human cells (Watanabe et al., 2019), and they facilitate the interaction of a CENP-C dimer with two CENP-A nucleosomes, at least *in vitro* (Walstein et al., 2021). A CDK1 phosphorylation site near the CENP-C motif enables high-affinity CENP-A binding, and ablation of this site renders human cells dependent on the CENP-C central domain (Ariyoshi *et al*., 2021; Watanabe *et al*., 2019). Like vertebrate CENP-C, yeast Mif2 is required for inner kinetochore assembly (Cohen et al., 2008). Its CENP-C motif binds Cse4/CENP-A and is required for viability (Cohen *et al*., 2008; Xiao et al., 2017), though a comparable CDK1-dependent step regulating nucleosome binding has not been described.

The extent to which cell cycle progression induces large-scale rearrangements of inner kinetochore sub-assemblies remains an open question. Evidence for such rearrangements has been detected in vertebrates (Ariyoshi *et al*., 2021; Hara and Fukagawa, 2020; Nagpal et al., 2015; Watanabe *et al*., 2019), motivating identification of the responsible post-translational protein modifications. What kinase activities are required, how might they influence inner kinetochore structure, and how does this relate to cell cycle progression?

To address these questions, we focused on the conserved and essential inner kinetochore protein, Mif2. Mif2 is heavily phosphorylated, the modified protein associates with assembled kinetochore particles, and phosphorylation accumulates in cells arrested at later stages of the cell cycle (S and metaphase) (Lang *et al*., 2018). We report here that Mif2 phosphorylation stabilizes the inner kinetochore. Our findings show an unexpectedly close relationship between this phosphorylation and components of the Ctf19c that are otherwise dispensable for viability. Overall, we connect cell cycle regulation to the functional plasticity of the inner kinetochore and discuss implications of this finding.

## Results

### Mif2 phosphorylation and interactions with Ctf19 complex factors

To better understand how Mif2 supports kinetochore assembly and stability, we used genetic complementation to identify the minimal Mif2 protein required for cell growth. Removal of Mif2 amino acid residues 1-200 (*mif2-Δ200*) supported viability, while further truncation to residue 256 (*mif2-Δ255*) did not (Figure 1A, Figure S2A). A finer deletion analysis covering the entire Mif2 N-terminal region (residues 1-240, Figure S1) yielded an internal deletion mutant that is hypersensitive to high temperature, hydroxyurea, and benomyl (*mif2-Δ181-240*, Figure S1). This part of Mif2 was previously named the PEST region (for <u>P</u>roline-, Glutamate (<u>E</u>)-, <u>S</u>erine-, and <u>T</u>hreonine-rich; Mif2-PEST), and temperature-sensitivity of *mif2-ΔPEST* cells is known (Cohen *et al*., 2008).

**Figure 1.**
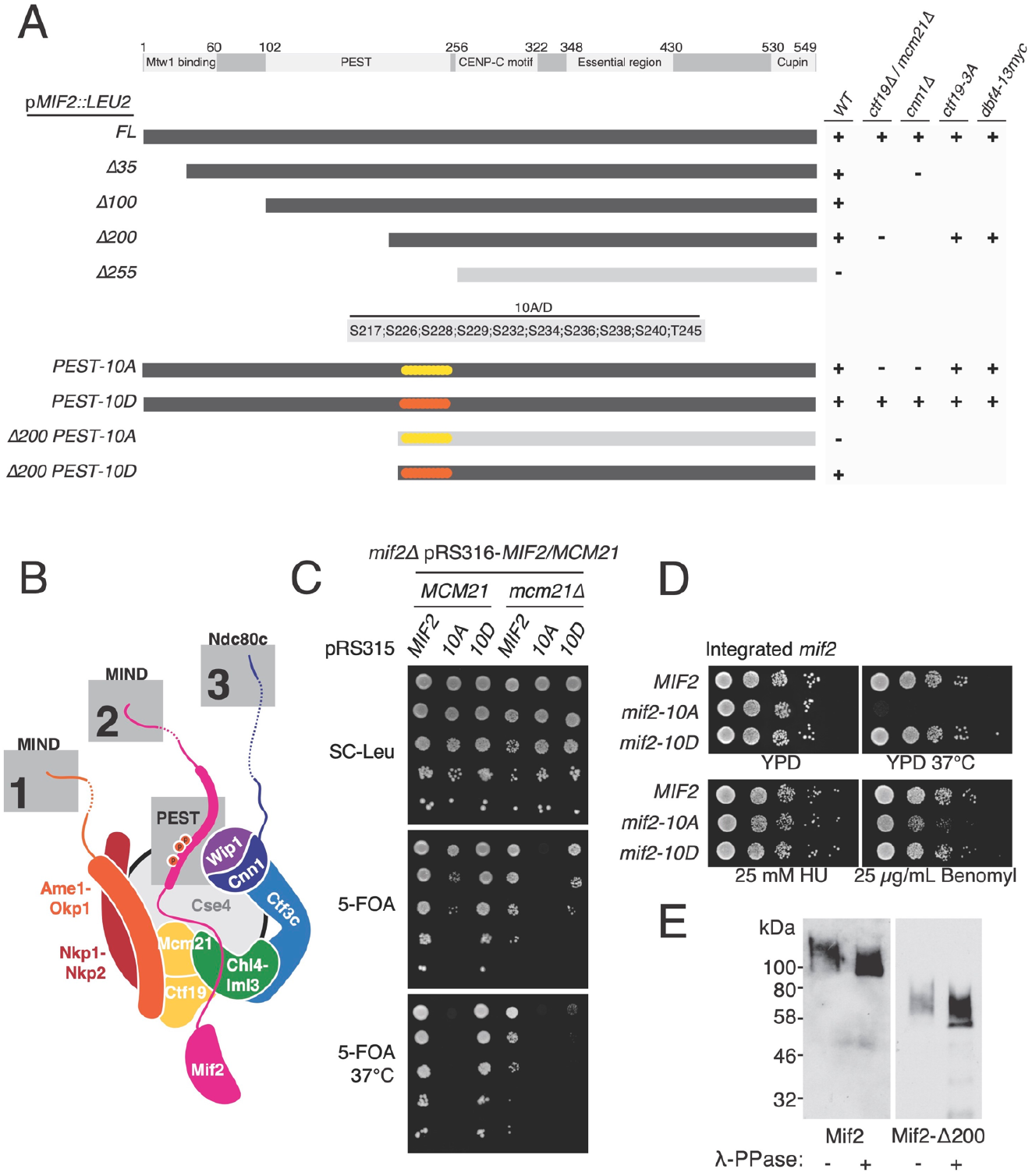
Mif2 PEST region mutations are lethal in Ctf19c mutant cells. A) Summary of genetic complementation experiments showing viability of the indicated *mif2* mutants (left). Viability of *MIF2-AID* cells with the indicated complementing *MIF2* plasmid was determined by serial dilution on plates with or without NAA (see also Figure S2). B) Illustration of the inner kinetochore and its outer kinetochore interaction motifs: 1. Ame1-N recruits MIND; 2. Mif2-N recruits MIND; 3. Cnn1-N recruits the Ndc80c. Disruption of motif 1 is lethal. Ctf19c deletion strains (e.g., *ctf19Δ, mcm21Δ*, and *cnn1Δ*) lack motif 3. *mif2-10A* mutations are in the PEST motif. C) Synthetic lethal interaction between *mcm21Δ* and *mif2-10A. mcm21Δ mif2Δ* double mutant cells carrying a complementing plasmid (*CEN/ARS-URA3* expressing *MCM21* and *MIF2*) and a test plasmid (*CEN/ARS-LEU2* expressing *MIF2, mif2-10A*, or *mif2-10D*) were plated on 5-FOA plates. D) Spot analysis to evaluate cell growth for the indicated genotypes exposed to heat stress (37 °C), replication stress (HU), or spindle stress (Benomyl). *MIF2, mif2-10A*, and *mif2-10D* were integrated into the native *MIF2* locus. E) Mif2 is phosphorylated in cells. Western blot (anti-FLAG) shows Mif2-3xFLAG electrophoretic mobility before and after phosphatase treatment (λ-PPase). Endogenously expressed Mif2-WT and Mif2-Δ200 were immuno-purified from asynchronous cultures (shorter exposure shown for Mif2-Δ200).

Characteristics of the Mif2-PEST region indicate it directs inner kinetochore assembly: it is immediately N-terminal to the Cse4-binding CENP-C motif, it is likely positioned near Ctf19c proteins in the assembled kinetochore, and its human counterpart binds CCAN factors (CENP-H/I, CENP-N) (Klare et al., 2015). Our unpublished biochemical reconstitution experiments indicated kinase activity is promotes robust inner kinetochore assembly. Therefore, we searched for putative Mif2-PEST phosphorylation sites and identified a cluster of 10 serine/threonine residues within this region (Figure 1, Figure S1). Mutation of these 10 residues to alanine was tolerated in full-length *MIF2* but lethal in *mif2-Δ200* (Figure 1A). Conversion of all 10 residues to aspartate in both full-length *MIF2* and *mif2-Δ200* supported cell viability.

Three well-characterized peptides recruit the microtubule-binding Ndc80 complex (Ndc80c) to centromeres in yeast (Figure 1B). These are Ame1-N, Mif2-N, and Cnn1-N (Dimitrova et al., 2016; Hornung et al., 2014; Malvezzi et al., 2013; Nishino et al., 2013; Przewloka et al., 2011; Screpanti et al., 2011; Thapa et al., 2015). Disruption of Ctf19c assembly in cells lacking Mif2-N is lethal (Killinger et al., 2020). Therefore, synthetic lethality between *mif2-10A* and *mif2-Δ200* suggests that a critically weakened kinetochore causes cell death. Indeed, the *mif2-10A* allele did not support the viability of *ctf19Δ* or *cnn1Δ* mutants, which have weakened kinetochores but are otherwise viable (Figure 1A, Figure S2). Neither the *ctf19-3A* mutation, which prevents Scc4-dependent centromeric cohesin recruitment (Hinshaw et al., 2017), nor the *dbf4-13myc* mutation, which prevents Ctf19c-dependent early replication of centromeric DNA (Natsume et al., 2013), recapitulated the viability defects seen in Ctf19c deletion strains expressing *mif2-10A*. Therefore, known Ctf19c functions not directly related to kinetochore-microtubule contact are dispensable for the viability of *mif2-10A* cells.

Extragenic *mif2-10A* caused a dominant defect in *mcm21Δ, ctf19Δ*, and *cnn1Δ* cells (Figure S2B). To circumvent this, we used a plasmid expressing both *MCM21* and *MIF2* to complement *mcm21Δ mif2Δ* cells and found that extragenic *mif2-10A* does not support viability in the *mcm21Δ mif2Δ* mutant background (Figure 1C). *mif2-10A*, when integrated in the *MIF2* chromosomal locus, supported cell growth, but the mutant cells were hyper-sensitive to elevated temperature and to benomyl (Figure 1D). *mif2-10D* showed normal growth in all conditions tested.

Kinetochore strengthening by Mif2 phosphorylation is the simplest explanation for the contrasting effects of *mif2-10A* and *mif2-10D*. Indeed, Mif2-Δ200 was phosphorylated when purified by means of a C-terminal 3-FLAG tag from *mif2-Δ200* cells (Figure 1E). Thus, in addition to previously described Ipl1-dependent N-terminal phosphorylation sites (Westermann *et al*., 2003), Mif2 is also phosphorylated at C-terminal sites. The genetic experiments described above indicate that phosphorylation sites between amino acid 200-250 likely control kinetochore activity.

### *DDK, Cdc5, and Ipl1 phosphorylate Mif2* in vitro

To determine the function of Mif2-PEST phosphorylation, we first identified the responsible kinases. The sequence context for known DDK and Cdc5 substrates matches the potential phosphorylation sites in the Mif2-PEST region (Botchkarev and Haber, 2018; Sheu and Stillman, 2006). We reconstituted DDK and Cdc5 kinase activities *in vitro* and tested Mif2 as a substrate. We also included Ipl1 in this analysis, because Mif2 is a known Ipl1 substrate (Westermann *et al*., 2003). Mif2 phosphorylation by DDK and Cdc5 has not been reported. All three kinases phosphorylated Mif2-WT (Figure 2A). Compared with Mif2-WT, the Mif2-10A protein served as a poor substrate for both Cdc5 and DDK (Figure 2B), indicating these enzymes target the residues mutated to make the Mif2-10A protein, at least *in vitro*. Cdc5 and DDK can act cooperatively; phosphorylation by one kinase primes further phosphorylation by the other (Princz et al., 2017). To test whether this is the case for Mif2, we compared Mif2-WT phosphorylation by each kinase alone to reactions containing both enzymes. Mif2-WT was phosphorylated to a greater degree by a mixture of Cdc5 and DDK than would be expected were both kinases working individually (Figure 2B). A Mif2 substrate lacking its N-terminal region recapitulated this effect (Mif2-Δ100, Fig. 2B). Therefore, Cdc5 and DDK cooperate to phosphorylate the Mif2 residues mutated in the Mif2-10A protein.

**Figure 2.**
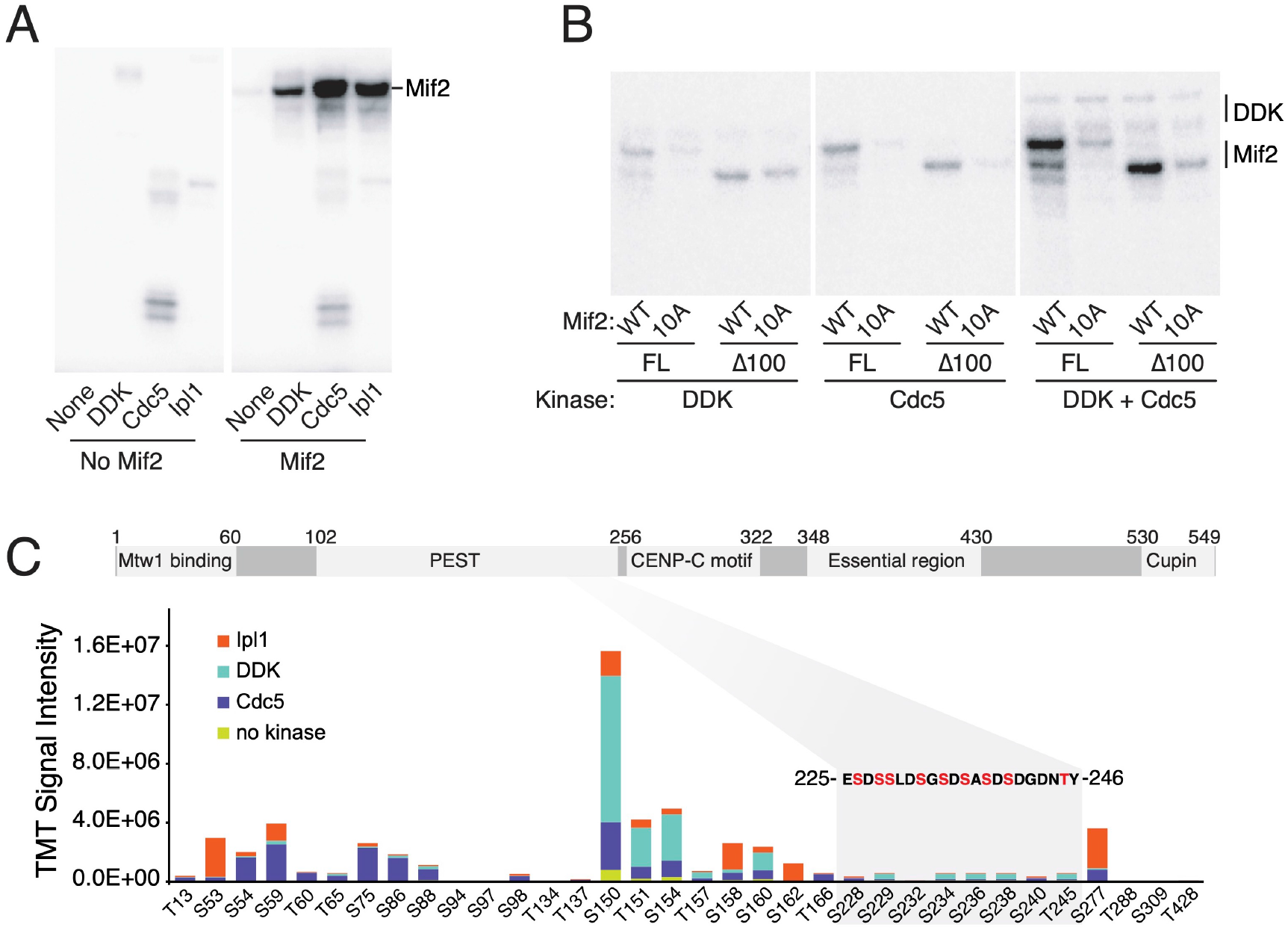
DDK and Cdc5 phosphorylate the Mif2 PEST region. A) Mif2 is a substrate of kinetochore-associated kinases DDK, Cdc5, and Ipl1. Purified Mif2 was incubated with the indicated kinases, and radioactive phosphate transfer was detected by autoradiography. B) Mif2-10A is a poor substrate for DDK and Cdc5. Kinase assays were performed as in panel A with the indicated substrates and enzymes (all panels from a single exposure). C) Mapping of Mif2 phosphorylation events *in vitro* by MS. Following *in vitro* phosphorylation of Mif2 by Cdc5, DDK, or Ipl1, phosphopeptides were TMT-labeled and quantified by MS (see Methods and Supplementary Table 1). Relative intensities of the same peptide treated with different kinases and unique TMT labels provide an accurate assessment of the relative contribution of each kinase to the abundance of a given phosphopeptide. Total signal intensities for different peptides primarily reflect unequal ionization efficiencies.

We used mass spectrometry (MS) to directly observe the products of Mif2 phosphorylation by Ipl1, Cdc5, and DDK. We applied Tandem Mass Tag (TMT) labeling to identify and quantify phosphorylated peptides after *in vitro* phosphorylation reactions (Figure 2C and Supplementary table 1). A given phosphopeptide’s propensity for ionization, combined with its abundance and level of phosphorylation, determines its detectability by MS. Thus, phosphopeptide detection by MS depends not only on the stoichiometry of phosphorylation, but also on the physical properties of the target species, making multi-phosphorylated peptides notoriously difficult to detect. Nevertheless, we found several phosphorylated peptides that mapped to the Mif2-PEST region after Mif2 treatment with Cdc5 and DDK, despite the fact the Mif2-PEST region is not well detected by this method. The Mif2 sites phosphorylated by DDK, Cdc5, and Ipl1 closely match the known consensus motifs of these kinases, as expected (Supplementary Table 1). MS analysis of Mif2-WT purified from yeast identified essentially the same phosphopeptides observed after *in vitro* kinase treatments (Supplementary Table 2), confirming that Cdc5, DDK, and Ipl1 are principally responsible for Mif2 phosphorylation in cells.

### Mif2 phosphorylation by Cdc5 and DDK regulates kinetochore assembly

Preferential recruitment of phosphorylated Mif2 to kinetochore particles assembled *in vitro* (Lang *et al*., 2018) suggests that the molecular function of Mif2 phosphorylation is to promote inner kinetochore assembly. To test whether Cdc5 and DDK enhance inner kinetochore assembly, we reconstituted this process using purified proteins and analyzed the resulting complexes by gel filtration chromatography. In the absence of kinase activity, Mif2-WT associated with the Ctf19c and the centromeric nucleosome, but overall complex formation was inefficient, a characteristic of this reconstitution that we have noted previously (Figure 3A, middle panel and our unpublished results). In contrast, Mif2-10D supported robust complex formation, consistent with a role for Cdc5 and DDK in promoting inner kinetochore assembly (Figure 3A, bottom panel).

**Figure 3.**
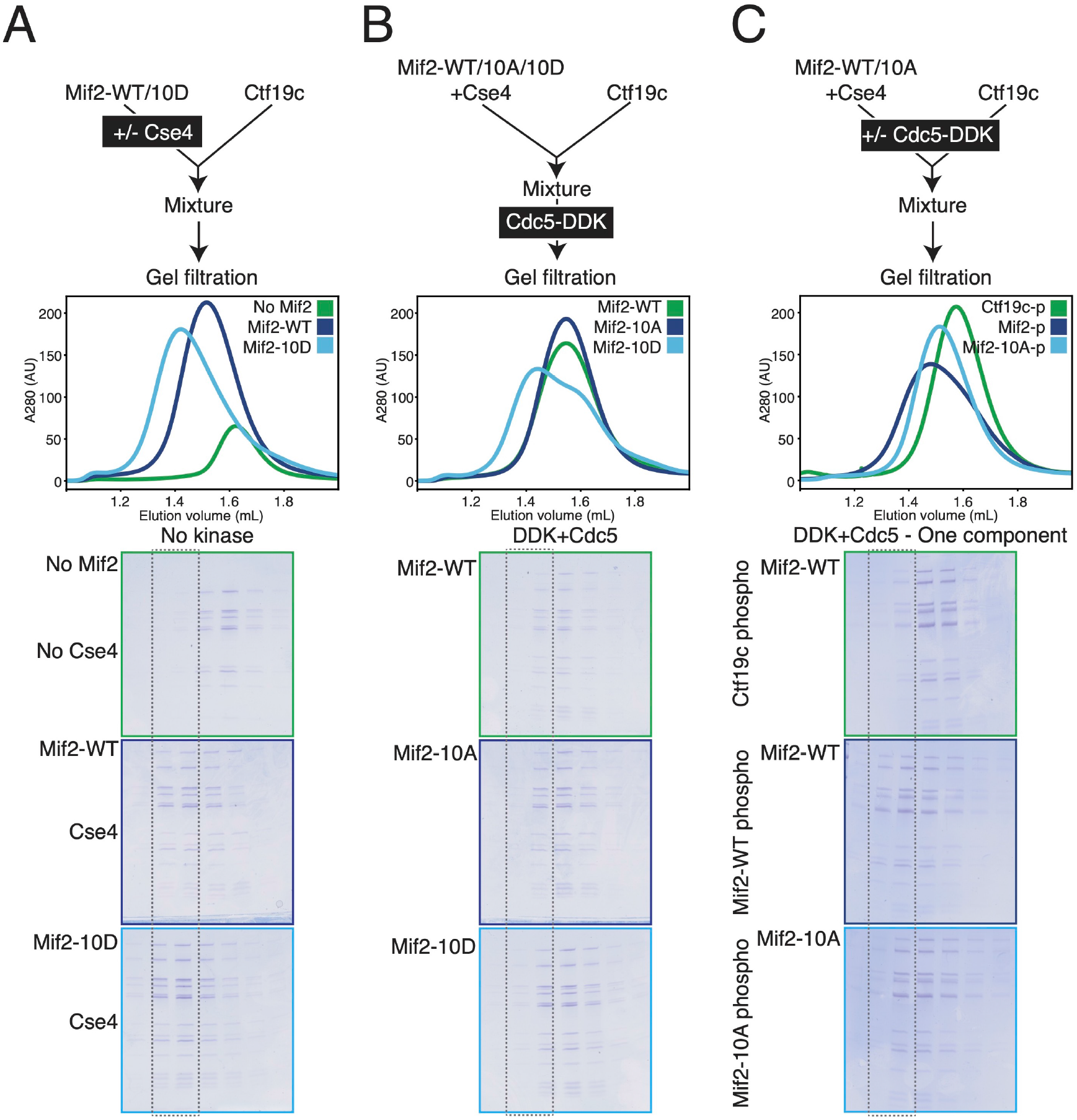
Biochemical reconstitution of inner kinetochore stabilization by DDK and Cdc5. A) The Mif2-10D-Cse4 nucleosome complex associates stably with the Ctf19c. The diagram shows experimental setup. Chromatogram shows the elution profile for the indicated samples (curve colors correspond to gel image outlines, dotted outline is a visual aid to highlight complex-containing fractions). All gel images show equivalent gel filtration fractions. B) DDK and Cdc5 treatment of preformed Mif2-Cse4 nucleosome-Ctf19c samples does not enhance inner kinetochore stability. Experiment performed as indicated in the diagram (top) and carried out as in panel A. C) DDK and Cdc5 treatment of the Mif2-Cse4 nucleosome complex enhances inner kinetochore assembly, and this requires Mif2-PEST phosphorylation. Either the Mif2-Cse4 nucleosome complex or the Ctf19c was phosphorylated before incubation with its counterpart as indicated in the diagram (top). The reactions were analyzed as in panels A and B. The phosphorylated components are indicated to the left of the gel images.

We next assessed the effects of Cdc5 and DDK activities on inner kinetochore formation using the same system. Kinase treatment of an assembly mixture containing all components before separation by gel filtration chromatography resulted in inefficient complex formation relative to the Mif2-10D sample not treated with kinases (Figure 3A, bottom panel versus Figure 3B, top panel). Thus, in addition to promoting inner kinetochore formation by phosphorylating Mif2, a second activity of Cdc5 and DDK inhibits complex formation in this system. To distinguish the opposing effects of kinase treatment, we treated either the Mif2-nucleosome complex or the Ctf19c with Cdc5 and DDK before the final incubation step (Fig. 3C, diagram). Treatment of the Mif2-nucleosome complex with kinases enhanced its interaction with the Ctf19c, and treatment of the Ctf19c had the opposite effect (Figure 3C). Mif2-10A could not be fully activated by kinase treatment. Therefore, Mif2 phosphorylation by Cdc5 and DDK is required for efficient inner kinetochore assembly, and a second activity of Cdc5 and DDK on the Ctf19c counteracts this effect, at least *in vitro* and in the absence of other kinetochore factors.

### Mif2 phosphorylation by Cdc5 and DDK enables Cse4-specific nucleosome binding

Mif2 binds Cse4 and promotes Ctf19c assembly (Westermann *et al*., 2003; Xiao *et al*., 2017). To determine how Mif2-PEST phosphorylation affects Cse4 binding, we used a gel shift assay (EMSA). In these experiments and those described above, Cse4 nucleosomes were wrapped with non-centromeric DNA, because nucleosomes wrapped with centromeric DNA could not be reliably produced *in vitro*. Mif2-10A bound a Cse4-contaning nucleosome with an affinity of ∼200 nM, while Mif2-10D showed a ∼10-fold lower affinity (Figure 4A). Wild type Mif2 bound the nucleosome with a similar affinity and migration pattern as Mif2-10A, and Mif2 phosphorylation by Cdc5 and DDK converted this high-affinity complex to one that resembled the Mif2-10D complex (Figure 4B). Mif2-WT showed a modest preference for Cse4-versus H3-containing nucleosomes (Figure S4A). This preference was abrogated by the 10A mutation and preserved by the 10D mutation. Mif2-10A has a higher affinity than Mif2-10D for the DNA used in these experiments (Figure S4B), explaining the observed differences in nucleosome affinities. Therefore, loss of DNA binding upon Mif2 phosphorylation enhances Mif2 selectivity for Cse4-over H3-containing nucleosome particles.

**Figure 4.**
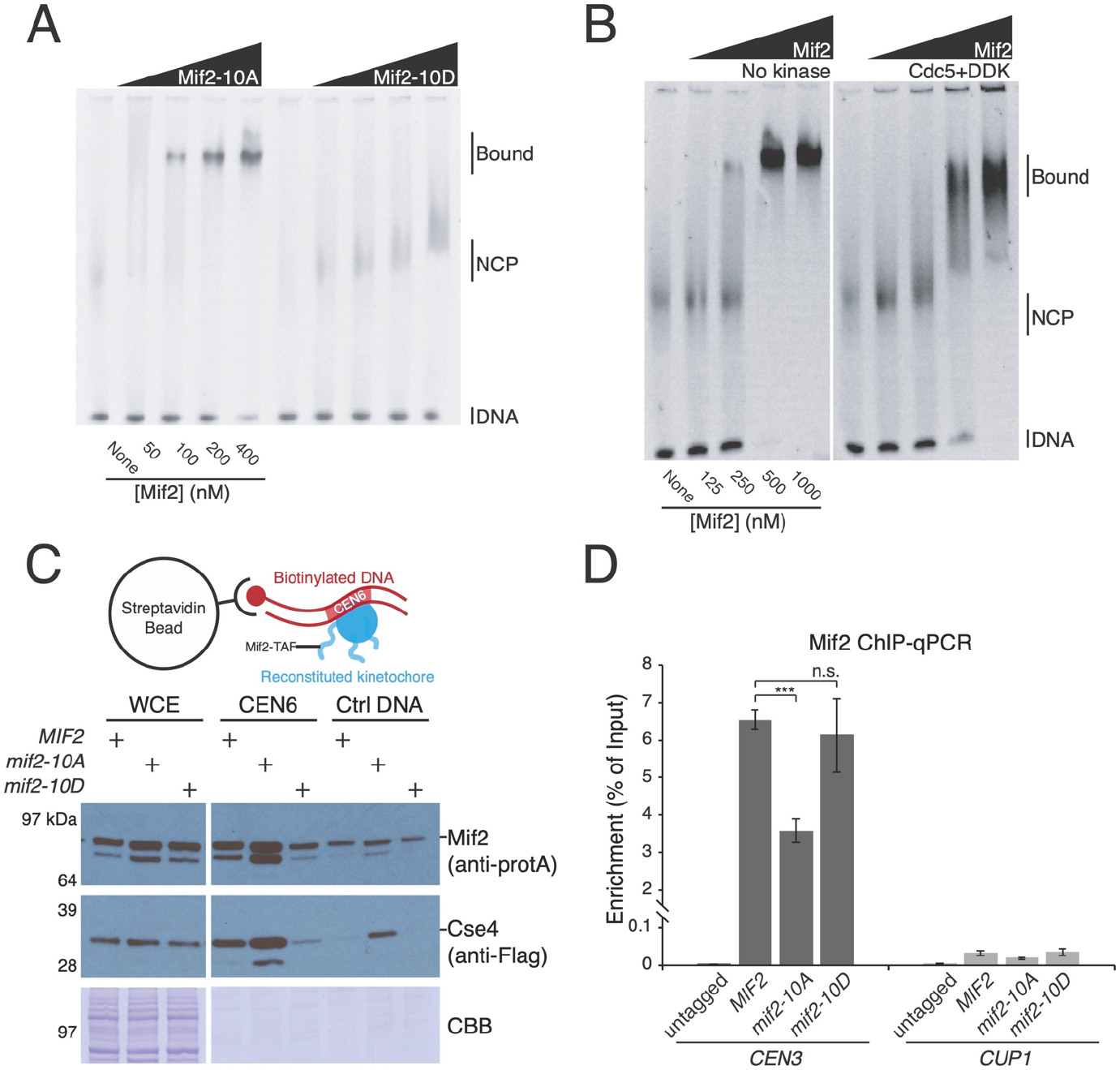
Mif2 phosphorylation modulates Cse4 nucleosome binding. A) Mif2-10A binds more tightly to the Cse4 nucleosome. The indicated Mif2 proteins were incubated with reconstituted Cse4-containing nucleosome samples, and the resulting complexes were analyzed by EMSA assays (Bound – Mif2-nucleosome complex; NCP – nucleosome core particle; DNA – free DNA). The Mif2 protein (above) and its monomeric concentration (below) are given. B) Mif2-PEST phosphorylation partially converts Mif2 to the low-affinity form seen in panel A. Mif2 was phosphorylated by DDK and Cdc5 before incubation with the Cse4 nucleosome and subsequent analysis by EMSA as in panel A. C) Mif2-centromere association in cell extracts recapitulates *in vitro* measurements (panel A). Mif2-WT, -10A, and -10D associate with centromeric DNA, and the association is decreased or unobservable for control DNA (Ctrl DNA; WCE – whole cell extract; CBB – Coomassie brilliant blue). CEN6 association is diminished for Mif2-10D. D) Mif2 phosphorylation enhances centromere binding *in vivo*. ChIP-qPCR experiments to measure Mif2-*CEN3* association were performed in cells with the indicated Mif2 mutations. Mif2 association with a non-centromeric locus (*CUP1*) is shown at right.

To extend these observations, we tested the association of cellular Mif2 and Cse4 with centromeric DNA, which also contributes to Mif2 recruitment (Xiao *et al*., 2017). Kinetochores assemble upon centromeric DNA incubated in cell extracts, and the reaction recapitulates *in vivo* assembly dependencies (Lang *et al*., 2018; Sorger et al., 1994). We incubated *CEN6* DNA with extracts made from asynchronous *MIF2, mif2-10A*, or *mif2-10D* cells and evaluated Mif2 and Cse4 recruitment by Western blotting (Figure 4C). Efficient recruitment depended on the centromere and matched the results of the EMSA experiments described above. Specifically, Mif2-*CEN6* association was stronger for Mif2-10A than for Mif2-10D. Mif2-10A, which displays higher DNA binding affinity *in vitro*, showed greater association with non-specific DNA, and this resulted in the co-recruitment of Cse4. Therefore, the Mif2-PEST mutations influence DNA affinity in extracts as they do for the purified proteins.

Lower overall affinity of Mif2-10D for the Cse4 nucleosome and for DNA indicates that inner kinetochore stabilization by Cdc5 and DDK requires cellular factors not included in the EMSA experiments and likely recruited at low efficiency (relative to Cse4 and Mif2) in extract assembly experiments. To test this, we performed chromatin immunoprecipitation and quantitative PCR experiments (ChIP-qPCR) to examine Mif2 localization. Mif2 localized to *CEN3* but not to a non-centromeric locus (Figure 4D). Mif2-10A was partially deficient in this localization, while Mif2-10D maintained robust *CEN3* association. Therefore, when measured *in vivo*, Mif2-centromere association does not strictly reflect the relative DNA binding efficiencies of the Mif2-10A and Mif2-10D proteins. The *in vivo* deficiency in centromere localization observed for Mif2-10A indicates that Mif2, once activated by Cdc5 and DDK, interacts with Ctf19c proteins to stabilize the inner kinetochore.

### *Mif2 phosphorylation is required for stable Mif2-Cse4 association* in vivo

To evaluate whether impaired Mif2-10A localization might correlate with a weakened Mif2-Cse4 association *in vivo*, we measured the interaction by co-immunoprecipitation. Cse4 co-purified with Mif2-WT, and less Cse4 co-purified with Mif2-10A (Figure 5A). Mif2-Cse4 binding was weakest in G1 and increased as cells progressed through the cell cycle. To directly compare Cse4 association with Mif2-WT and Mif2-10A, we performed competitive binding experiments by overexpressing an extragenic copy of Mif2 (Figure 5B). Overexpressed Mif2-WT but not Mif2-10A interfered with the ability of endogenous Mif2-WT to bind Cse4 (Figure 5C). Therefore, Mif2-10A is defective in its association with Cse4 *in vivo*. These results, considered together with the findings presented above, indicate that Mif2 phosphorylation by Cdc5 and DDK stabilizes the inner kinetochore by converting Mif2 from a high-affinity DNA binder to a bridging factor that specifically and stably binds Cse4 to promote kinetochore assembly.

**Figure 5.**
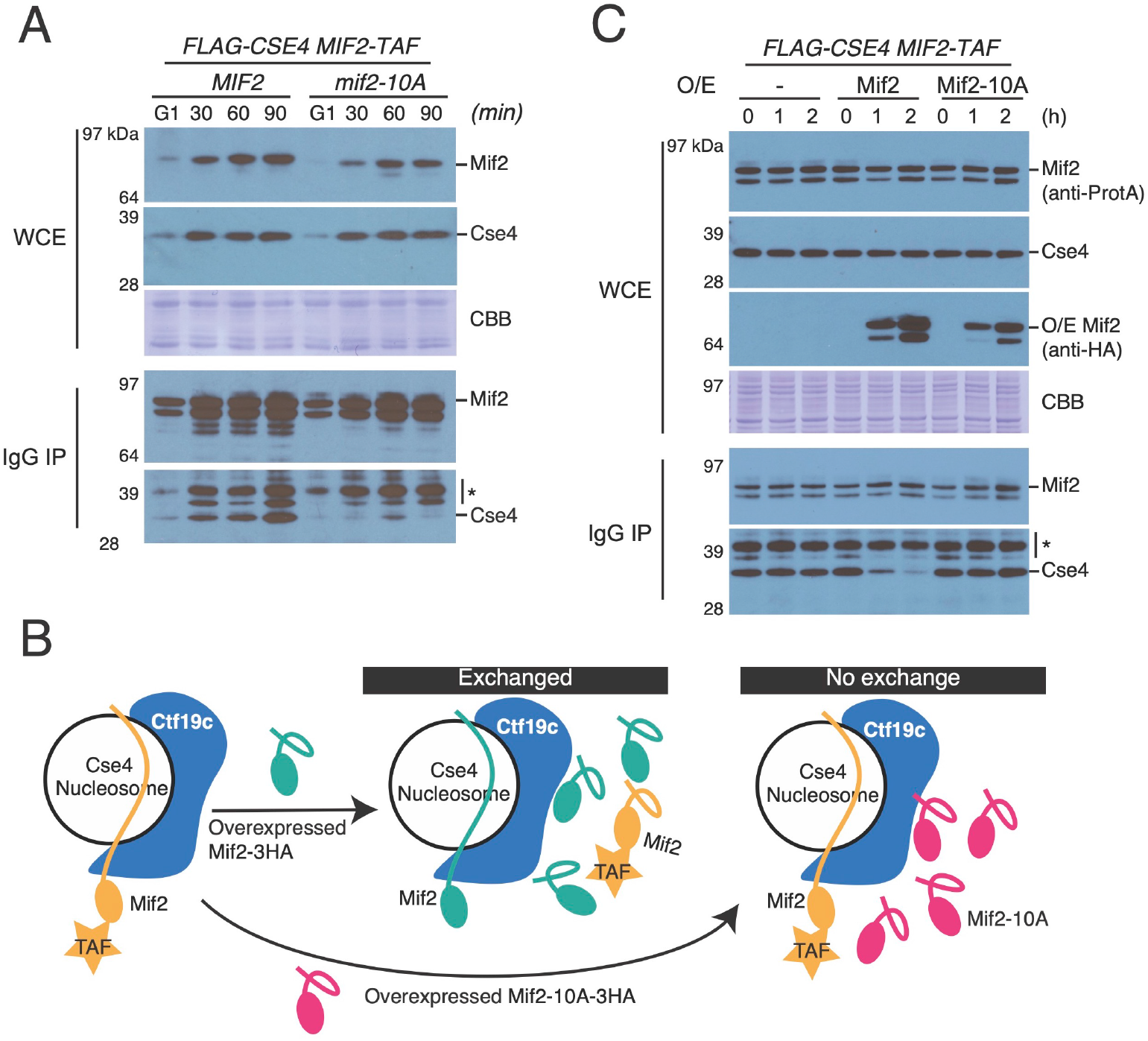
Mif2 phosphorylation is required for stable Mif2-Cse4 interaction *in vivo*. A) The *mif2-10A* mutation prevents stable Mif2-Cse4 association. Cells were arrested in G1 with alpha factor and released into the cell cycle for the indicated amounts of time (top). Mif2-TAF was immunopurified and co-purifying FLAG-Cse4 was detected by Western blotting (WCE – whole cell extract; CBB – Coomassie brilliant blue). B) Schematic showing the experimental design for competition pull-down experiments (panel C). Endogenously expressed Mif2-TAF at the centromere is challenged by induced expression of the Mif2-3HA (WT or 10A mutant; IgG IP – Mif2-TAF immunoprecipitation). C) Overexpressed Mif2 but not Mif2-10A interferes with endogenous Mif2-Cse4 association. Pulldown experiments were performed as in panel A except time points were taken after induction of extragenic Mif2-WT or Mif2-10A. Endogenous Mif2-TAF was purified and Cse4 association was detected by Western blotting (O/E – overexpression).

Extragenic *mif2-10A* shows partial dominance over *MIF2* in Ctf19c-deficient cells. To test whether Ctf19c factors safeguard the kinetochore by preventing the deleterious effects of Mif2-10A incorporation, we carried out Mif2 competition experiments like those described above. The Cse4-Mif2-10A interaction is not stabilized in *mcm21Δ* cells, ruling out a protective function for Ctf19c factors in the face of *mif2-10A* expression (Figure S5). Therefore, the Ctf19c dependence of *mif2-10A* cells implies that *mif2-10A mcm21Δ* (for example) kinetochores are weaker than those of either individual mutant and that they do not support chromosome segregation.

### Mif2 phosphorylation and Ctf19c assembly cooperate to ensure kinetochore stability

Overall, our findings show that robust kinetochore assembly depends on Mif2-PEST phosphorylation by Cdc5 and DDK and that this phosphorylation enhances inner kinetochore assembly. As a qualitative measure of kinetochore function *in vivo*, we examined the ability of cells to tolerate a dicentric plasmid (Figure 6A). Mitotic cells establish biorientation for each centromere independently. A single dicentric chromatid can attach to opposite spindle pole bodies, mimicking the merotelic attachments seen in organisms with multivalent centromeres (Dewar et al., 2004). Budding yeast have no mechanism to correct this error, and the result is a severe viability defect (Koshland et al., 1987). We confirmed that deletion of non-essential Ctf19c factors restores viability to cells bearing a dicentric plasmid (Poddar et al., 1999). The *mif2-10A* but not the *mif2-10D* mutation also rescued viability, indicating kinetochore disfunction in the mutant cells is a consequence of weak and not aberrantly strong attachments. The shared phenotype between cells lacking Ctf19c proteins and those deficient in Mif2 phosphorylation indicates the presence of parallel pathways for stable microtubule-kinetochore attachment.

**Figure 6.**
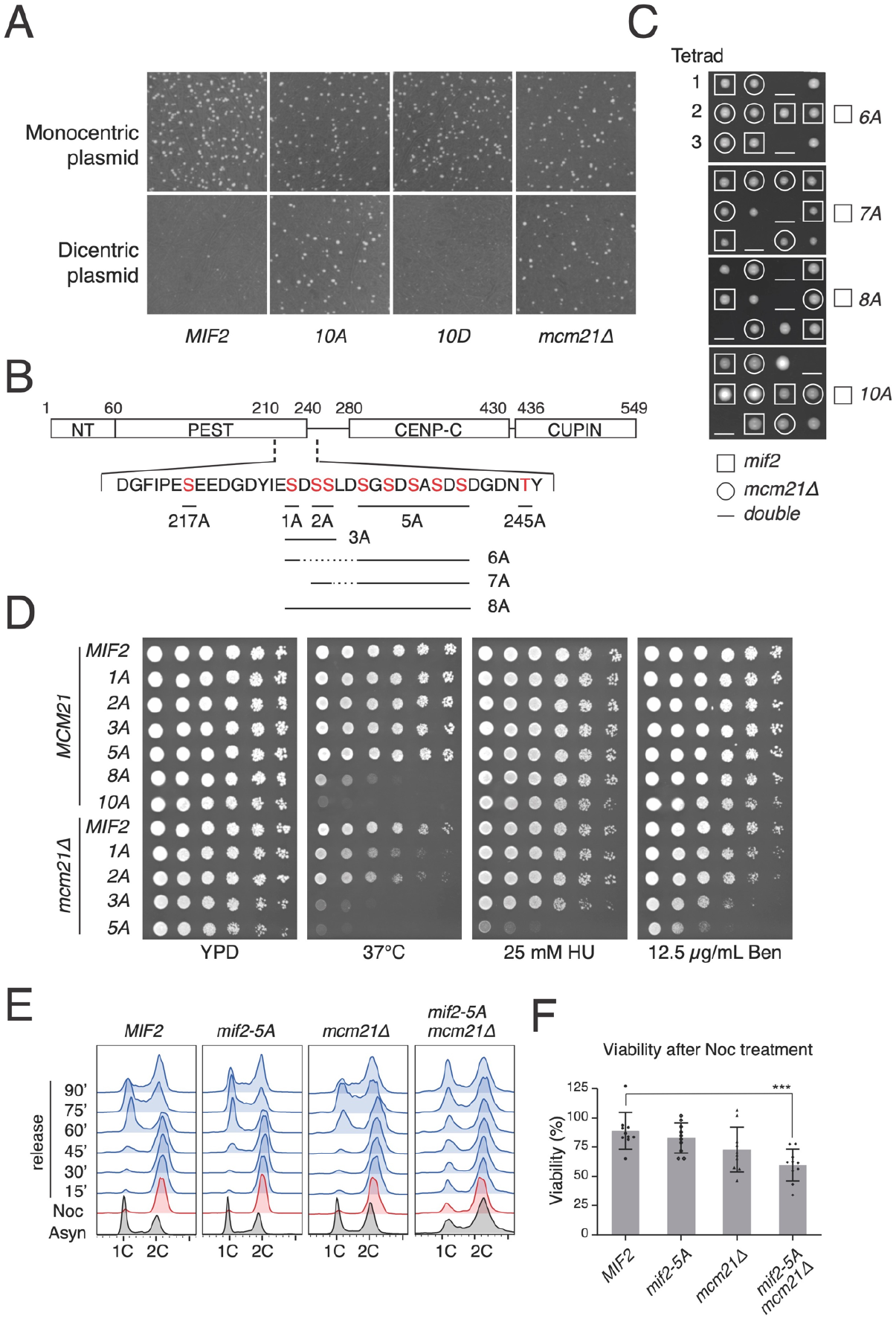
Cooperative Mif2 phosphorylation supports kinetochore function. A) *mif2-10A* cells have impaired kinetochore function. Strains with the indicated genotypes were propagated with a monocentric or dicentric plasmid as indicated. Viability of transformants under selection for the plasmid was assessed by colony formation. B) Diagram showing the Mif2-PEST region and the phosphorylated residues mutated to make the *mif2-1A, -2A, -3A, -5A, -6A, -7A*, and *-8A* mutations. C) Multiple phosphorylation site mutations cause lethality in combination with *mcm21Δ*. Heterozygous diploid strains (*MCM21/mcm21Δ*; *MIF2/mif2-6A, -7A, -8A*, or -*10A*) were sporulated, and the meiotic products were genotyped as indicated. D) Removal of Mif2 phosphorylation sites causes increasing sensitivity to heat stress (37 °C), replication stress (HU), and spindle stress (Ben). E) Cell cycle analysis shows that *mif2-5A mcm21Δ* cells fail to recover from a mitotic arrest. Cells of the indicated genotypes were arrested in G2/M by nocodazole treatment for 3 hours and released into fresh medium. Reentry into the cell cycle is indicated by the reappearance and subsequent disappearance of the 1C peak at 60 minutes and 90 minutes, respectively. F) *mif2-5A mcm21Δ* cells show a significant loss of viability upon release from a nocodazole arrest.

To further examine the functional requirement for Mif2 phosphorylation, we created a series of *MIF2* alleles coding for differing amounts of phospho-null mutations (Figure 6B). Sporulation of heterozygous *mcm21Δ mif2-10A* cells confirmed that *mif2-10A* cells are viable but depend on an intact Ctf19c (Figure 6C). Likewise, *mif2-8A, -7A*, and *-6A* mutant cells depend on *MCM21* for viability. In contrast, we recovered viable spores for *mif2-1A, -2A, -3A*, and *-5A mcm21Δ* double-mutant cells, and these had increasing sensitivities to heat stress (37 °C), hydroxyurea (HU), and benomyl (Ben) (Figure 6D). The *mif2-3A* and *mif2-5A* mutations ablate non-overlapping subsets of phosphorylation sites. Together, these residues make up the *mif2-8A* mutation that renders cells dependent on *MCM21* for viability. The nearly identical phenotypes of the *mif2-3A* and *mif2-5A* mutants indicates there is not a necessary or sufficient phosphorylation site in the Mif2-PEST region. Instead, the *MIF2* allelic series strongly suggests that kinetochore stabilization depends on multiple Mif2-PEST phosphorylation sites, consistent with the biochemical DDK/Cdc5 cooperativity described above.

To determine how the *mcm21Δ mif2-5A* double mutant cells respond to acute mitotic stress, we arrested cells in metaphase with the spindle poison nocodazole and examined cell cycle progression upon release from this treatment (Figure 6E). In response to nocodazole, wild type cells arrested in G2 and synchronously re-entered the cell cycle upon release. *mcm21Δ* and *mif2-5A* cells responded similarly, indicating that neither mutation is sufficient to completely destabilize the kinetochore. In contrast, *mcm21Δ mif2-5A* double mutant cells arrested in G2 but showed evidence of defective chromosome segregation upon release (Figure 6E, right column). This was most apparent 90 minutes after release, when the double mutant cultures were essentially asynchronous, indicating long and variable delays in the execution of mitosis. Accumulation of G2/M cells in unsynchronized *mcm21Δ mif2-5A* cultures supports this interpretation (Figure 6E, bottom row). We also measured cell viability, normalized to the beginning of the procedure, of cells released from a nocodazole arrest (Figure 6F). Whereas wild type, *mcm21Δ*, and *mif2-5A* cells largely maintained their viability throughout the arrest, the viability of *mcm21Δ mif2-5A* cells dropped significantly. Therefore, the kinetochores of *mcm21Δ mif2-5A* cells do not generate productive microtubule attachments upon spindle regrowth.

## Discussion

Kinetochores must accommodate the events of the cell cycle. These include the replication of underlying centromeric DNA, the subsequent reattachment to spindle microtubules, the establishment of tension-sensing end-on microtubule attachments, and the maintenance of end-on attachments during anaphase. Post-translational modifications of kinetochore proteins accompany the transitions between these functions. We have investigated this conserved property of kinetochores by studying Mif2, a central regulator of kinetochore activity that integrates cell cycle cues. Mif2 phosphorylation by Cdc5 and DDK enhances inner kinetochore assembly. Defective Mif2 phosphorylation renders cells inviable when the Ctf19c is compromised. The findings provide evidence for an inner kinetochore assembly mechanism that depends on coordinated post-translational modification of a basal component (Figure 7).

**Figure 7.**
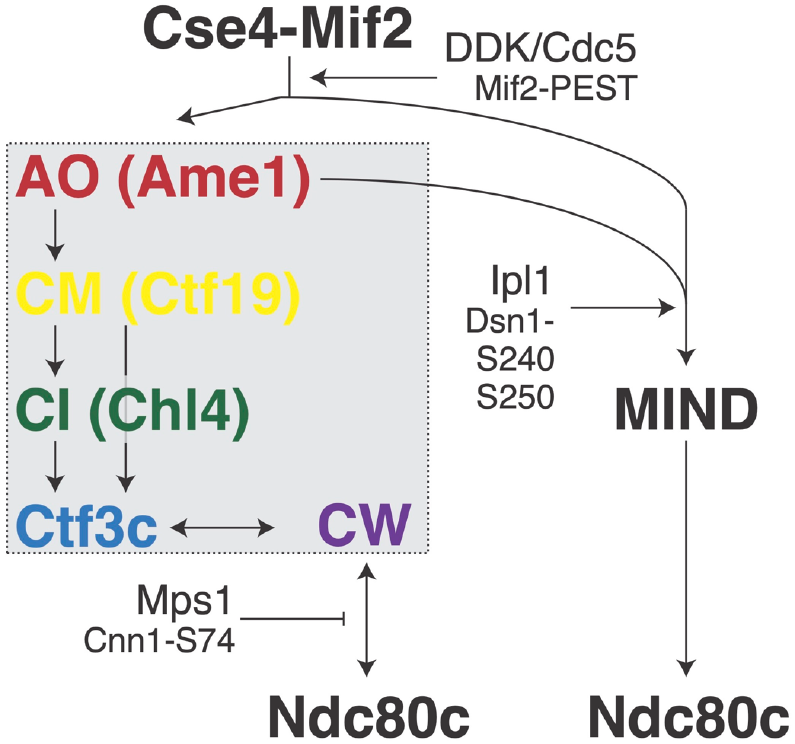
Inner kinetochore assembly mechanism. Diagram showing known interactions and kinase activities supporting inner kinetochore assembly (AO – Ame1-Okp1; CM – Ctf19-Mcm21; CI – Chl4-Iml3; Ctf3c – Ctf3-Mcm16-Mcm22; CW – Cnn1-Wip1). Factors higher in the diagram support recruitment of those beneath. The gray box demarcates the Ctf19c.

How might Cdc5 and DDK prepare the inner kinetochore for mitosis? DDK phosphorylates the inner kinetochore just before centromere-proximal origin firing, which marks the onset of S phase (Hinshaw *et al*., 2017; Natsume *et al*., 2013). Biochemical association of DDK with Cse4 begins in S phase and persists through mitosis (Mishra et al., 2021), while Cdc5 associates with Cse4 in G2/M phase (Mishra et al., 2016; Mishra et al., 2019). It is not known exactly when Cdc5 phosphorylates its many kinetochore substrates (Klemm et al., 2020; Lera et al., 2016; Olafsson and Thorpe, 2020). We favor a model in which DDK phosphorylates Mif2 at the G1/S transition, and this primes Mif2 for subsequent phosphorylation by Cdc5. If so, Mif2-dependent inner kinetochore strengthening may only occur after a given centromere has activated its associated origin, preventing premature strengthening of a kinetochore built upon an unreplicated centromere. The extensive Mif2 phosphorylation *in vivo* has so far prevented direct observation of PEST phosphorylation in Mif2-WT, making the development of a tool for this purpose a priority to study this model further.

Upon activating Mif2 for kinetochore assembly, Cdc5 and DDK reduce its affinity for the Cse4/CENP-A nucleosome, at least *in vitro* and in cell extracts (Figure 4). Two non-exclusive mechanisms could explain this switch from DNA binding to inner kinetochore stabilization. First, Mif2-PEST phosphorylation might convert Mif2 from a DNA binder to a Ctf19c binder. Second, Mif2 phosphorylation might liberate a nucleosomal binding site for the Ctf19c. Mif2 autoinhibition by a fold-back mechanism has been proposed (Killinger *et al*., 2020), and a CDK-dependent fold-back has been observed in a cryo-EM structure of chicken CENP-C bound to a CENP-A nucleosome (Ariyoshi *et al*., 2021). Our results imply that Mif2-PEST phosphorylation inhibits DNA binding by the Mif2 AT-hook, as suggested by Brown (Brown, 1995) and as has been observed for the DNA binding domain of the Ets-1 transcription factor (Pufall et al., 2005; Serber and Ferrell, 2007). Structural and biochemical studies are required to evaluate these possibilities.

The modified Mif2-PEST residues are not obviously conserved at the primary sequence level. Nevertheless, many or all Mif2/CENP-C homologues contain an analogous PEST region (Walstein *et al*., 2021). Clustered phosphorylation sites are common features of proteins with cell cycle-dependent functions (Nash et al., 2001; O’Neill et al., 1996; Serber and Ferrell, 2007). Sic1 is an archetypal example; clustered phosphorylation sites create a switch-like response to rising kinase activity, rendering the protein susceptible to Cdc4-dependent degradation (Nash et al., 2001). The phosphorylation sites in the Mif2-PEST region share properties with Sic1-N (multiple sites and poor primary sequence conservation), indicating that they also constitute a nonlinear switch that responds to kinase activity. This might be crucial for coordinating centromere-specific biochemical states among all 32 kinetochores. The graded cellular response to successively inactivated *MIF2* alleles (*mif2-10A, -8A, -5A*, etc.) strongly supports this functional model for Mif2-PEST phosphorylation.

In addition to the positive feedback described above, our study of Mif2-PEST phosphorylation has shown that two systems-level positive feedback loops converge on Mif2 to regulate kinetochore assembly. The first involves cooperative activation of Mif2 by Cdc5 and DDK. The DDK step necessarily occurs before inner kinetochore stabilization and therefore in the absence of robust Ctf3 localization, which is required for maximal DDK localization (Hinshaw *et al*., 2017). Indeed, kinetochore-associated Ctf3-GFP intensity is at its nadir as cells approach S-phase (Hinshaw and Harrison, 2020). Thus, DDK-dependent Mif2-PEST phosphorylation produces more robust Ctf19c assembly, which in turn results in further Mif2-PEST phosphorylation.

Cdc5/PLK1 localization depends on CENP-U binding in vertebrates (Kang et al., 2006; Singh et al., 2021) and likely also in yeast (Ame1/Okp1-N-Cdc5 phosphorylation, our unpublished data). Thus, Cdc5, like DDK, positively regulates its own kinetochore activity. Dual mutually reinforcing positive feedback loops ensure robust Mif2 activation and consequent inner kinetochore stabilization.

The findings we have presented lead us to three main conclusions. The simplest conclusion is that Mif2 phosphorylation stabilizes the inner kinetochore. The second conclusion is that robust kinetochore assembly involves feed-forward loops that progressively stabilize key interfaces. We have found one such example involving clustered phosphorylation sites in Mif2. Other clustered sites probably exist in the kinetochores of yeast and other organisms, enabling switch-like responses to noisy input signals. The third conclusion is that integration of multiple cell cycle signals by the inner kinetochore produces an assembly mechanism that is robust to perturbation. This is seen from the lethal effect of combining *mif2-10A* with mutations affecting the Ctf19c. The kinase activities required for chromosome segregation rise and fall at rates that depend on nutrient signaling and physical forces. Perturbation of any individual signal might yield a temporary defect that can be overcome by positive feedback, a property that ensures robust coupling between kinetochore assembly and cell cycle progression.

## Acknowledgments

We thank Stephen Harrison and Kevin Corbett for helpful discussions and for comments on the manuscript. We thank Phong Lee for help with nucleosome reconstitutions. This work was supported by NIH GM116897, OD023498, and University of California CRCC faculty seed grant to H.Z. S.M.H. was supported in part by funding from HHMI and the Helen Hay Whitney Foundation.

---

**Figure S1.**
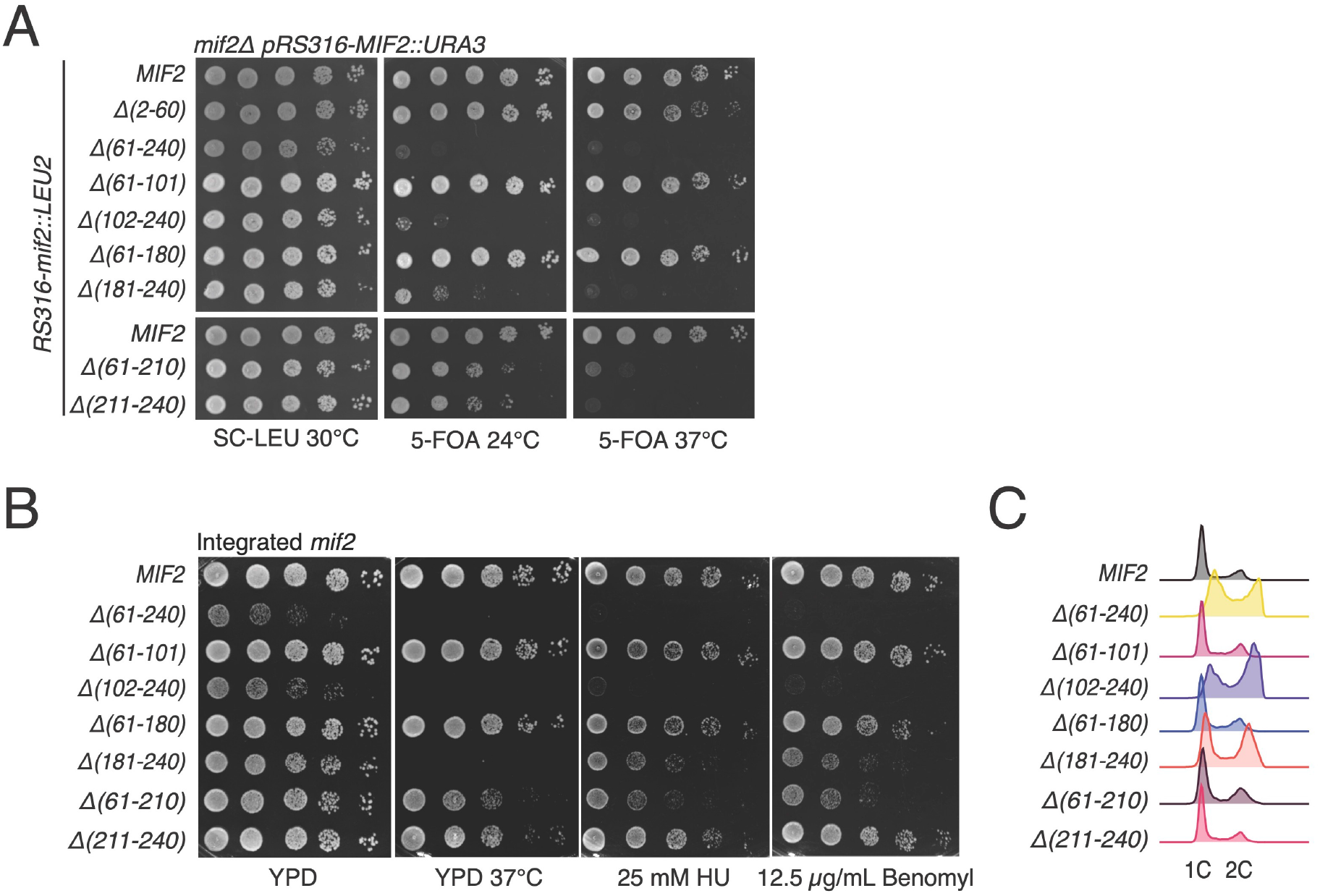
Deletion analysis of the Mif2-PEST region and phenotypic analysis of associated mutants. A) The *mif2-Δ181-240* mutant does not support normal viability. A plasmid-shuffling assay was used to measure the ability of the indicated *mif2* alleles to support viability (*mif2Δ; pRS316-MIF2::URA; pRS315-mif2::LEU2*). Numbers correspond to deleted amino acid residues in the expressed proteins. Neither *mif2-Δ61-210* nor *mif2-Δ211-240* completely phenocopies *mif2-Δ181-240*, indicating the presence of functional elements on both the N- and C-terminal sides of Mif2 residue 210. B) Viability analysis as in panel A was repeated with the indicated *mif2* alleles integrated into the *MIF2* locus. Stress conditions are indicated beneath images. C) FACS analysis showed that *mif2* mutants with more severe growth impairment tend to accumulate more G2-DNA content.

**Figure S2.**
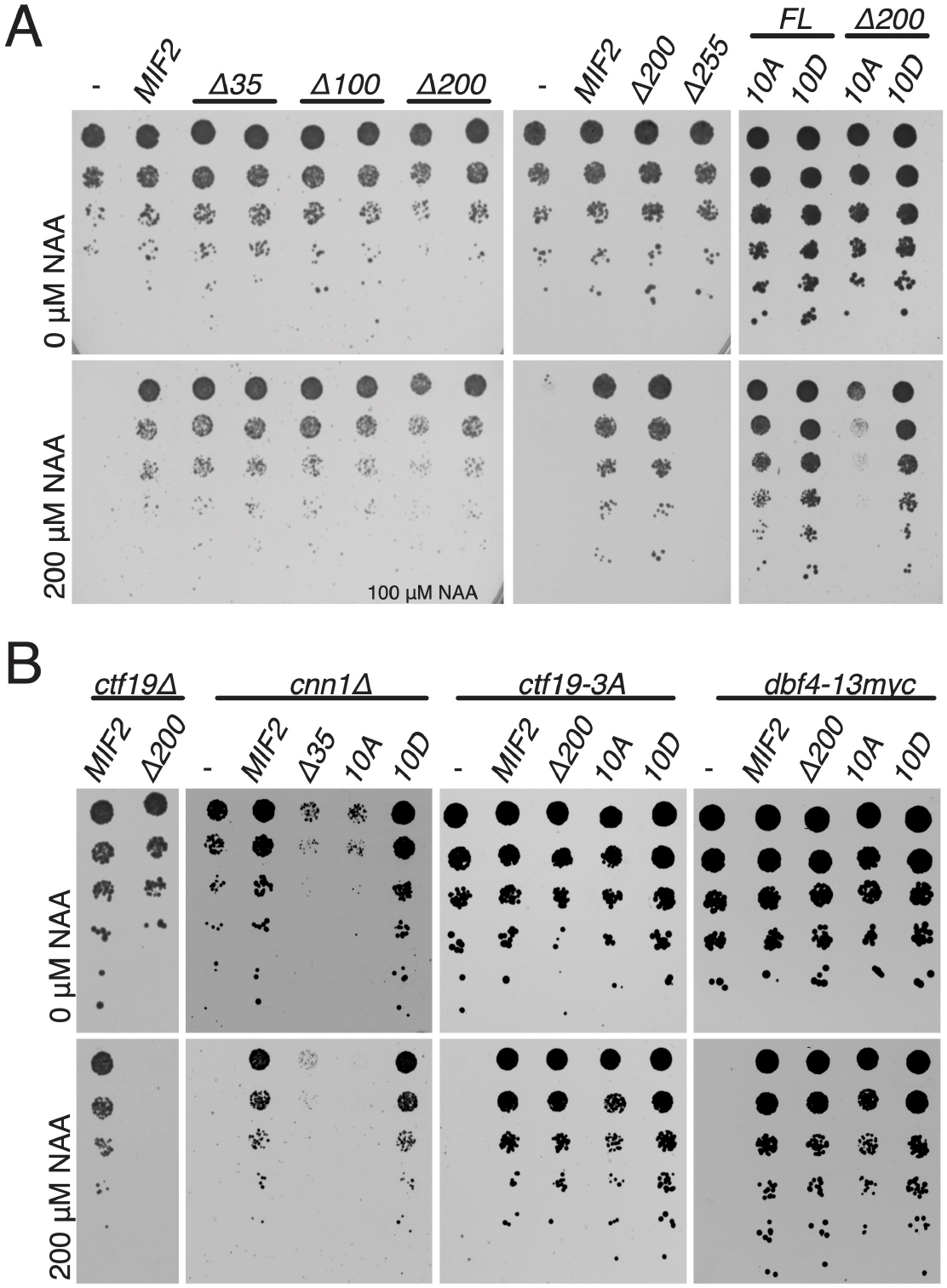
Mif2 genetic complementation tests and analysis of associated mutants. A) Identification of a minimal C-terminal Mif2 fragment and identification of the Mif2-10A mutant. Endogenous Mif2 was depleted by auxin treatment (NAA – 1-Napthaleneacetic acid), and the indicated *MIF2* alleles were supplied on a centromeric plasmid coding for *MIF2* and ∼500 bp flanking chromosomal DNA. B) Complementation analysis as in panel A shows that *CTF19* and *CNN1* but not cohesin recruitment (*ctf19-3A*) or early centromere replication (*dbf4-13myc*) are required for viability in cells expressing *mif2-ΔN* or *mif2-10A*.

**Figure S4.**
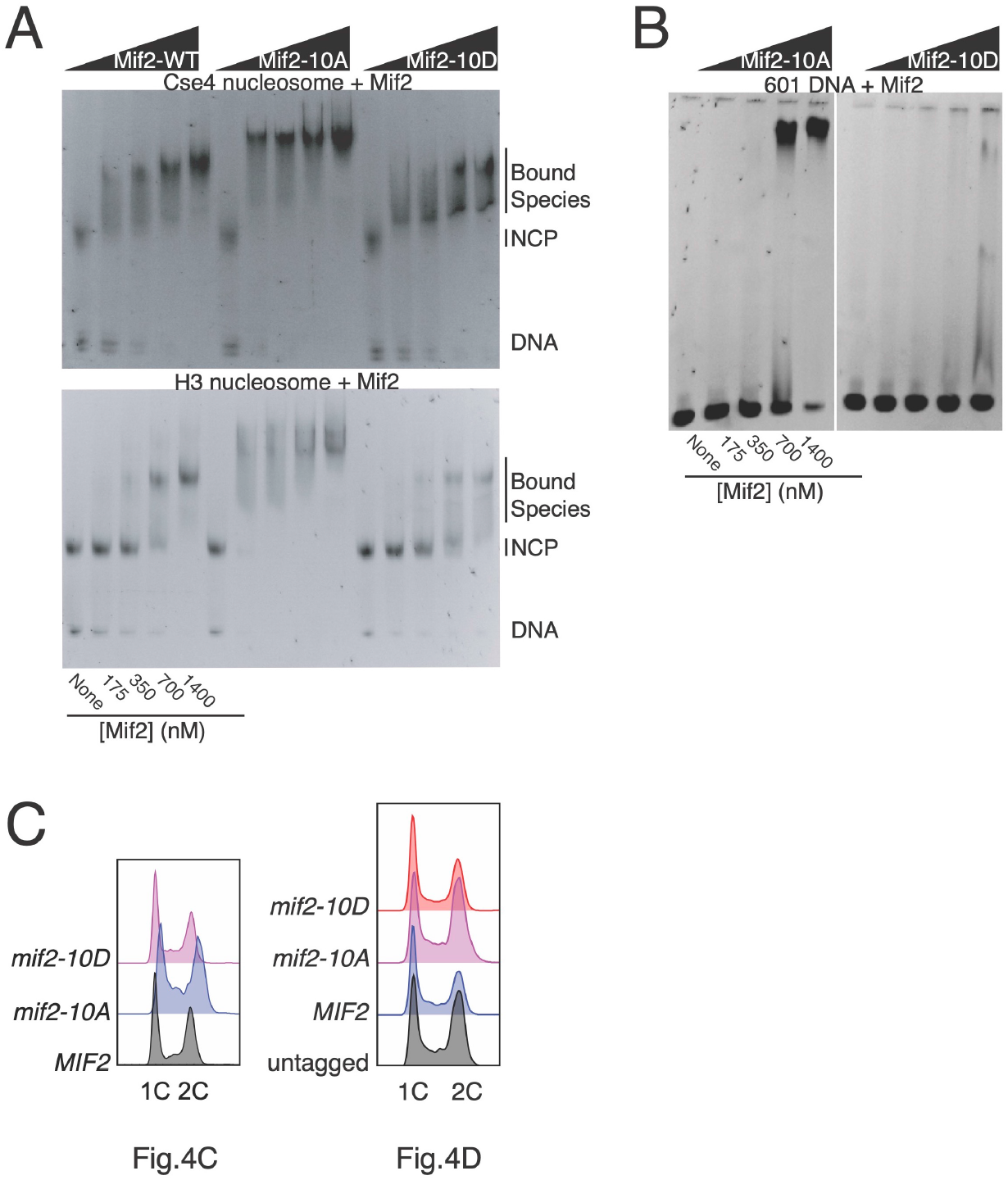
Selectivity and DNA binding of Mif2-WT, -10A, and -10D. A) Mif2 selectively binds Cse4-versus H3-containing nucleosomes. EMSA was performed as in Figure 4A-B. B) Mif2-10D DNA binding is not detectable. C) Cell cycle profiles for cultures used in Figure 4C and Figure 4D as indicated.

**Figure S5.**
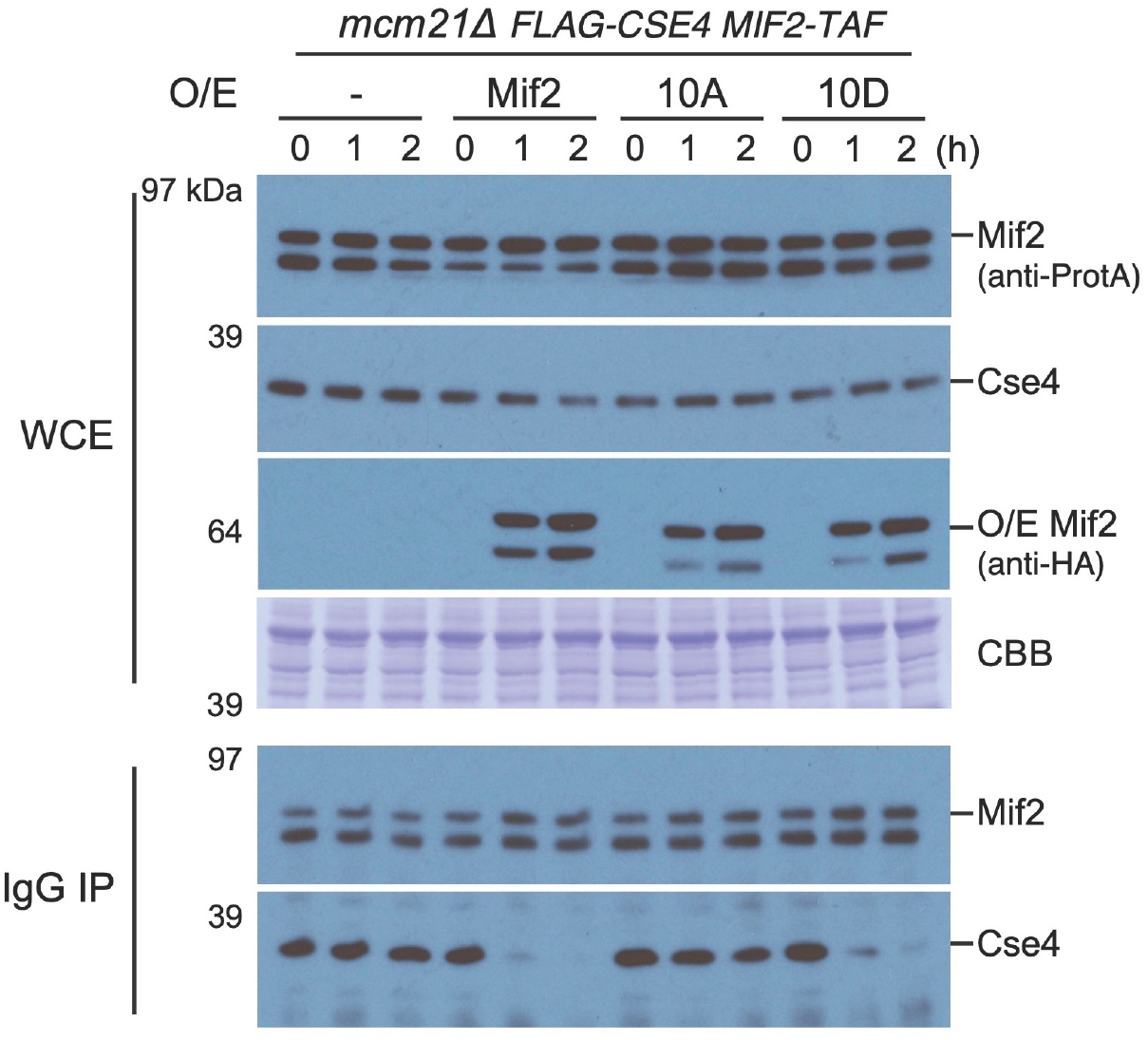
Mcm21 does not safeguard against deleterious Mif2-10A incorporation, and Mif2-10D competes with Mif2-WT. Mif2 competition pulldown experiments were performed as in Figure 5C, except the strain background was *mcm21Δ*.

## Methods

### Key Resources Table

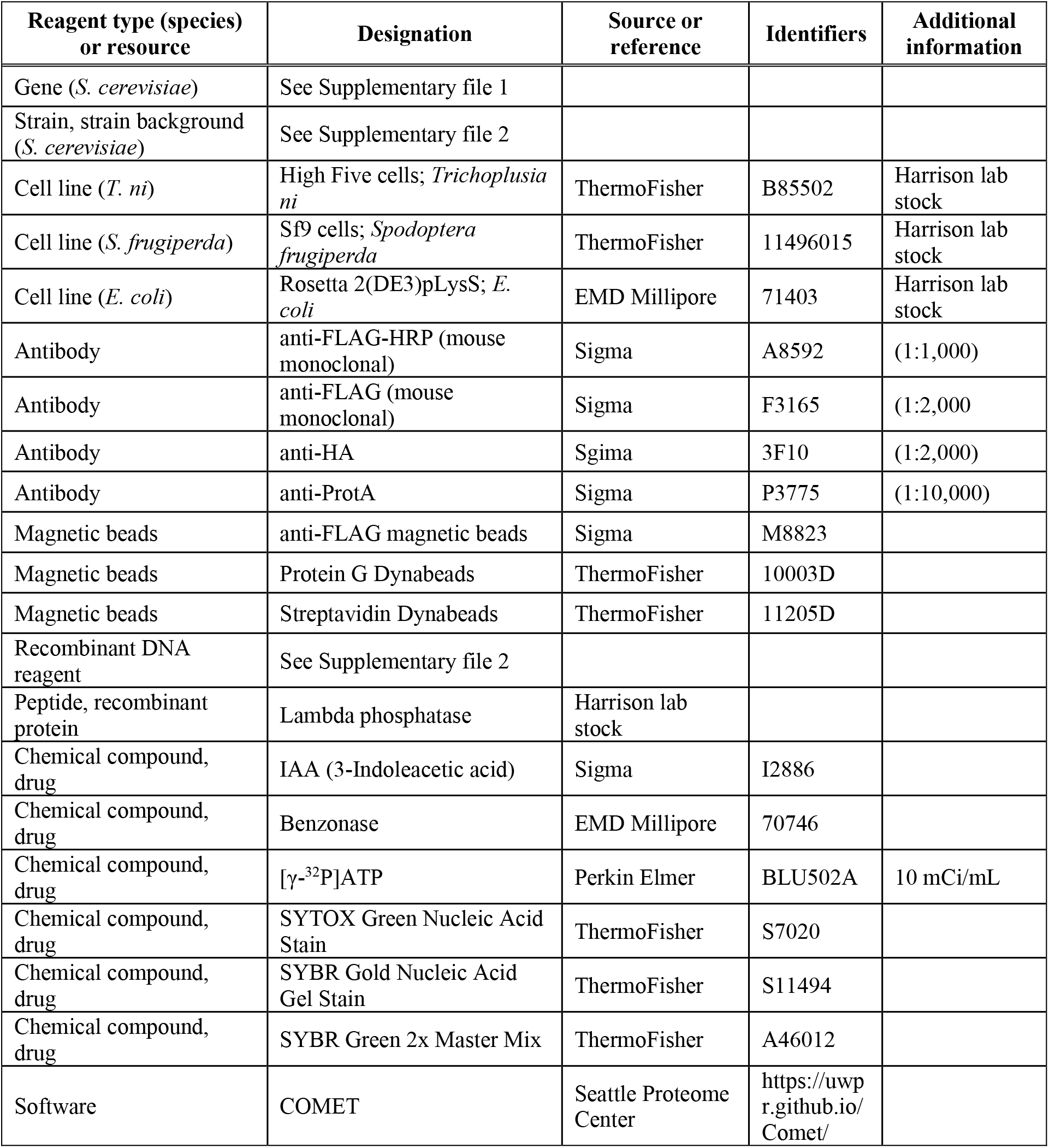

### Yeast plasmid and strain construction and growth conditions

A genomic fragment of Mif2 (chrXI:292980-275333) was cloned in CEN/ARS plasmids (pRS315 and pRS316) via yeast recombination repair. To generate *MIF2-TAF::HisMX6* (TAF: 6xHIS-3xFLAG-ProteinA), the Mif2-expressing plasmid was linearized by SphI and a PCR product of TAF:HisMX6 was fused to *MIF2* C-terminus via yeast recombination. Deletion and point mutations of Mif2 were then introduced in these plasmids (Supplementary Table 3).

Restriction fragments of plasmids containing various *mif2* mutants, selected by His-dropout, were integrated into the chromosomal *MIF2* locus in the strain with its genomic *mif2Δ* mutation complemented by the *CEN/ARS*-bearing plasmid pRS316*-MIF2*. His-positive colonies were replicated on plates containing 5-fluoroorotic acid (5-FOA) to evict the complementing pRS316*-MIF2*. Successful integration was confirmed by nourseothricin-sensitivity and PCR tests. Transformations for genomic modifications and plasmid introductions were done by standard methods (Longtine et al., 1998). All strains were grown at 30 °C in yeast extract peptone dextrose medium (YPD) unless otherwise indicated.

### Viability tests for Mif2 complementation experiments

Multiple methods were used to determine whether the described *mif2* mutants support cell growth. For *MIF2-AID* experiments, PCR integration was used to tag endogenous *MIF2* in a strain expressing *TIR1* (*pADH1-OsTIR1-9xMyc*). Centromere plasmids coding for *MIF2* alleles specified (including ∼500 bp flanking genomic sequence on either side) were subsequently transformed and selected for using the *LEU2* marker. Transformants were picked, streaked, and grown overnight in synthetic complete medium without leucine (SC-leu) before plating a five-fold dilution series. For strains with modifications at the endogenous *MIF2* locus, cells were grown and plated on the indicated media as for *MIF2-AID* cells.

### Mif2-FLAG pull-down and phosphatase treatment

Asynchronously dividing cultures were grown in YPD and harvested by centrifugation. Cell pellets were washed once in lysis buffer (25 mM HEPES, pH 7.5; 2 mM MgCl_2_; 0.1 mM EDTA; 0.5 mM EGTA; 0.1 % NP-40; 175 mM potassium glutamate; 15 % glycerol by volume; 2 μg/ml aprotinin, pepstatin, and leupeptin; 1 mM PMSF and benzamidine) and frozen at -80 °C. Cell pellets were thawed by addition of lysis buffer supplemented with phosphatase inhibitors (80 mM NaF; 20 mM NaVO4; 80 mM β-glycerol phosphate) and lysed by bead beating. Cleared supernatants were mixed with anti-FLAG magnetic beads and incubated at 4 °C for one hour with end-over-end nutation. Beads were washed four times with lysis buffer containing phosphatase inhibitors before a final wash in lysis buffer supplemented with 10 mM MnCl_2_ and without phosphatase inhibitors. Beads were incubated in MnCl_2_-containing buffer for ten minutes at 30 °C with or without excess lambda phosphatase before removal of unbound material and heating with SDS-PAGE sample buffer. Anti-FLAG Western blotting was used to visualize immunopurified Mif2 (anti-FLAG-HRP; Sigma A8592).

### Cell cycle analysis by flow cytometry

Liquid cell culture at each desired time point was fixed with 70 % ethanol (final concentration). All samples were left in ethanol at -20 °C overnight. The next day, fixed cells were collected by centrifugation, resuspended in 50 mM sodium citrate pH 7.0 with RNaseA (250 μg per sample) and Proteinase K (250 μg per sample), and incubated overnight at 37 °C. On the third day, the cells were collected by centrifugation, resuspended in 50mM sodium citrate with 1 μM Sytox Green (ThermoFisher), and sonicated at 80 kHz for 3-5 seconds. After incubation in a dark room for an hour, the samples were analyzed, and data was collected with the BD LSRFortessa X-20 cytometer.

### Recombinant protein expression and purification

Ipl1 kinase was expressed in *E. coli* Rosetta 2(DE3) cells (EMD Millipore). A polycistronic co-expression vector coding for Sli15-580-698 and Ipl1-AS6 (Kung et al., 2005) was constructed by ligation independent cloning (LIC) and overlapping PCR. Both proteins had N-terminal 6-His tags. Cdc5 was cloned by LIC into a vector coding for an N-terminal 6-His-MBP tag. Histone proteins were cloned by LIC and overlapping PCR to create co-expression plasmids coding for all four histone proteins on a single transcript. Histone H2A and codon-optimized histone H4 carried N-terminal 6-His tags. DDK (Dbf4 and Cdc7-AS3; (Wan et al., 2006)) was cloned into a single baculovirus transfer vector by LIC (Gradia et al., 2017). Both proteins had N-terminal 6-His tags. Ctf19c components were cloned sequentially into BioBrick-enabled LIC vectors (Gradia et al., 2017) to create two coexpression plasmids coding for eleven of the thirteen components (Ame1-Okp1-Nkp1-Nkp2-Ctf19-Mcm21 and Ctf3-Mcm16-Mcm22-Cnn1-Wip1). The remaining two Ctf19c components, Chl4 and Iml3, were purified from *E. coli* transformed with pSMH104 (Hinshaw and Harrison, 2013). Mif2 and associated mutants were cloned into a baculovirus transfer vector using LIC.

For protein expression in *E. coli*, cells were grown at 37 °C to OD ∼1 before induction with 0.4 mM isopropyl β-D-1-thiogalactopyranoside (IPTG). Cells were incubated overnight at 18 °C before harvesting by centrifugation and resuspension in buffer D800 (20 mM HEPES, pH 7.5; 10 mM imidazole, pH 8.0; 150 mM NaCl; 10 % glycerol by volume; 2 mM β-mercaptoethanol) supplemented with 2 μg/ml aprotinin, leupeptin, and pepstatin and 1 mM PMSF and benzamidine. Cells were stored at -80 °C until purification. For protein expression in insect cells, High Five cells (*T. ni*, ThermoFisher) were grown in EX-CELL 405 medium (Sigma) and infected with P3 virus at a cell density of ∼1 million/mL. 72 hours after infection, cells were harvested by centrifugation, resuspended in buffer B50 (B800 with 50 mM NaCl) supplemented with protease inhibitors and stored at -80 °C until purification. Baculovirus stocks were generated according to the manufacturer’s recommendation in Sf9 cells (*S. frugiperda*, ThermoFisher).

For protein purification from both *E. coli* and insect cells, lysis was done by sonication, and cell extracts were clarified by centrifugation at 43549.6 ×*g* for 60 minutes. The NaCl concentration was adjusted to 800 mM or 2 M before lysis for insect cell or histone octamer expressions, respectively. 6xHis-tagged proteins were purified by Co^2+^ affinity and eluted in C150 (D800 with 400 mM imidazole and 150 mM NaCl; Cdc5), C100 (D800 with 100 mM NaCl and 400 mM imidazole; DDK, Ipl1-Sli15, Ctf19c components, Mif2 and mutants), or C1000 (D800 with 1M NaCl, 400 mM imidazole, and 10 mM EDTA; histone octamers).

For kinases, the following further purifications were carried out. Ipl1-Sli15 and DDK were applied to a 5 mL cation exchange column (HiTrap SP HP; GE) and eluted with a linear gradient from B50 (Ipl1-Sli15) or B100 (DDK) to D800. Peak fractions were pooled, concentrated by ultrafiltration, and applied to an S200 column (S200 10/300; GE) equilibrated with GF150 (20 mM Tris-HCl, pH 8.5; 150 mM NaCl; 1 mM TCEP). Cdc5 was concentrated by ultrafiltration before injection onto an S200 column equilibrated with GF150. The pooled eluate was concentrated by ultrafiltration, glycerol was added to a final concentration of 5% by volume, and protein was stored at -80 °C until use.

For Ctf19c reconstitution, the Ame1-Okp1-Nkp1-Nkp2-Ctf19-Mcm21 and Ctf3-Mcm16-Mcm22-Cnn1-Wip1 complexes were purified separately as described above. Eluates from the metal affinity step were applied to a 5 mL cation exchange column (HiTrap SP HP; GE). Peak fractions were pooled, concentrated, and mixed at an equimolar ratio at a final volume of ∼1 mL, to which a molar excess of TEV-cleaved Chl4-Iml3 was added. Lambda phosphatase, MnCl_2_ (final concentration 1 mM), and benzonase (∼10 U/mL final concentration) were added, and the reaction was incubated at 30 °C for one hour before application to an S200 column equilibrated with GF150. The pooled eluate was concentrated and stored as described above.

Mif2 and associated mutant proteins were purified exactly as for Ctf19c factors with the following exceptions. A 5 mL anion exchange column (HiTrap Q HP; GE) was used. Peak fractions were treated with lambda phosphatase but not benzonase. Concentrated and dephosphorylated Mif2 was applied to an S200 column equilibrated in GF500 (GF150 with 500 mM NaCl) before concentration and freezing as described above.

Histone octamers were further purified by concentration and application to an S200 column equilibrated in GF1000 (GF150 with 1000 mM NaCl). Recombinant nucleosome core particles were reconstituted on 147 bp 601 DNA by salt gradient dialysis. 601 DNA was amplified by PCR from a plasmid template using the following oligonucleotides at large scale:

oSMH1950 – ATCGAGAATCCCGGTGCC
oSMH1951 – ATCGGATGTATATATCTGACACGTGC.

Reaction products were pooled, diluted with water and 100 mM HEPES, pH 7.5, 10 mM EDTA, and 400 mM NaCl (final concentrations) before purification by anion exchange chromatography (HiTrap Q HP; GE). Peak fractions were precipitated with 70 % ethanol by volume (final) at -20 °C, pelleted, reconstituted in 50 mM Tris-HCl, pH 8.5, and stored at -20 °C until use. For nucleosome wrapping reactions, a 1.2 molar ratio of histone octamer to 601 DNA was mixed in NCP-hi (1000 mM NaCl, 10 mM Tris-HCl pH 7, 1 mM EDTA, 1 mM dithiothreitol (DTT)), and overnight gradient dialysis was carried out at room temperature into NCP-lo (NCP-hi with 10 mM NaCl). After a final ∼2 h dialysis in NCP-lo, the sample was analyzed by native gel electrophoresis as described below. Nucleosome particles were used for EMSA or reconstitution experiments within 48 hours of gradient dialysis.

### In vitro *kinase assays*

The indicated kinases and substrates were mixed before addition of an equal volume of ATP/[γ-^32^P]-ATP (0.2 mM ATP-MgCl_2_, 0.5 μC/μL [γ-^32^P]ATP) in 2x kinase buffer (100 mM HEPES, pH 7.5; 10 mM Mg(OAc)_2_, 20% (v/v) glycerol, 400 mM potassium glutamate, 2 mM EDTA, 0.02% (v/v) NP-40 substitute; 4 mM NaoVO_4_; 40 mM NaF; 2 mM β-mercaptoethanol) to start the reaction. Reactions were incubated at 30 °C for one hour before heating with SDS-PAGE sample buffer. Reaction products were separated by SDS-PAGE and dried gels were imaged using autoradiography.

### TMT-mass spectrometry to measure phosphopeptide abundance

To analyze the purified and kinase-treated Mif2 proteins, we applied Tandem Mass Tag (TMT) labeling system (ThermoFisher) to identify and quantify phospho-peptides within each kinase-treated sample (Ipl1, DDK, Cdc5, or none). Briefly, 15 μg purified Mif2 proteins, with or without kinase treatment, were first dissolved in 50 μL of 6 M urea with 50 mM NH_4_HCO_3_. Then, each sample was treated with 10 mM DTT to disrupt the disulfide bonds and subsequently alkylated with 30 mM iodoacetamide. 1 μL beta-mercaptoethanol was added after 30 minutes to quench the alkylation reaction. 5 μL of the quenched samples were diluted in 25 μL of 50 mM NH_4_HCO_3_ and digested by trypsin or chymotrypsin for 2 hours at 37 °C. To end the digestion, trifluoroacetic acid (TFA) was added to a final concentration of 0.2 % in each sample. Each set of samples, digested by trypsin or chymotrypsin, was combined, desalted by C18 columns, and dried completely before resuspension in 20 μL of 50 mM KH_2_PO_4_ (pH 8.0). Each sample was treated with 8 μL of TMT labeling reagent at room temperature overnight. The next day, 1 μL of 1 M Tris-HCl, pH 8.0 was added to quench any unincorporated labeling reagent. After labeling, all samples were combined, desalted by C18 columns, and dried under vacuum. The dried sample was resuspended in 0.5 % acetic acid and processed for analysis using a ThermoFisher Orbitrap Fusion LUMOS Tribrid mass spectrometer as described (Suhandynata et al., 2019).

To identify the *in vivo* phosphorylation sites of Mif2, 1 liter culture of Mif2-TAF cells grown in YPD at OD_600_ ∼ 1.0 were collected and lysed with a PBS buffer containing protease inhibitors, PMSF, 0.5 M NaF, 0.5 M beta-glycerophosphate, 20 mM EDTA, and 1 μM Okadaic acid. Cells were lysed by glass beads beating at 4 °C for up to 2 hours (1 minute breaking, 2 minutes cooling program). After centrifugation, the clarified lysate was collected and incubated with 100 μL of anti-FLAG M2 beads (Sigma) at 4 °C overnight. The next day, anti-FLAG beads were washed with 1 mL of ice-cold lysis buffer for 5 times and eluted in 2 quick steps by 300 μL 0.1 M Glycine-HCl (pH 2.0). The total elution volume was 600 uL and each elution was completed within 15 seconds. After evaluation of Mif2 purification by Western blot (anti-Protein A), the samples were neutralized, reduced, alkylated, digested with trypsin, acidified, and desalted using the protocol described above for the *in vitro* phosphorylated samples.

To enrich phosphopeptides, the treated sample was purified with an IMAC (immobilized metal affinity chromatography) column as described (Suhandynata et al., 2014). Briefly, to make fresh IMAC columns, the beads were recovered from a Qiagen Ni-NTA spin column (Cat No. 31014, one spin column is enough to make ∼70 μL IMAC beads) and Ni^2+^ were stripped by gently shaking in 50 mM EDTA, 1 M NaCl for an hour, then washed with ddH_2_O, 0.6 % acetic acid, and recharged with Fe^3+^ by gently rotating in 50 mL of 0.1 M FeCl_3_ in 0.3 % acetic acid for an hour. Beads were washed once with 0.6 % acetic acid, then twice with 0.1 % acetic acid. Once prepared, beads can be stored in 0.1 % acetic acid for up to a week at 4 °C. To purify phosphopeptides, we packed 10 μL of IMAC beads in a gel loading tip, resuspended dried peptides in 40 μl of 1 % acetic acid (pH 3-4), then loaded the peptide sample slowly into the IMAC column by slightly applying syringe pressure. The IMAC column was washed twice with one bed volume of 0.6% acetic acid and once with a half bed volume of ddH_2_O. Phosphopeptides were eluted with three bed volume of 6 % NH_4_OH, dried, and resuspended in 5 μL of 0.6 % acetic acid for MS analysis, as described above. COMET (Seattle Proteome Center: Trans Proteomic Pipeline) software package was used for database searching. A static mass modification of 57.021464 Da for cysteine residues and a differential modification of 79.966331 Da for Ser/Thr phosphorylation were used.

### Inner kinetochore reconstitution experiments

The indicated components were mixed at equimolar concentrations at a final volume of 50 μL. For kinase treatments, the relevant complexes were mixed with purified kinases and ATP-MgCl_2_ (1 mM final concentration) and incubated for 1 hour at 30 °C. Buffer and nucleotide were added to unphosphorylated reactions, which were also incubated at 30 °C. Subsequent binding reactions were performed on ice for one hour in the presence of 2 mM ADP (final concentration) before separation on a Superose 6 column (5/150 GL) equilibrated in GF150-HEPES (GF150 with HEPES, pH 8.6 replacing Tris-HCl). Identical fractions were collected and analyzed by SDS-PAGE for all experiments.

### Gel shift assay (EMSA)

Recombinant Mif2 and nucleosome samples were mixed in a final volume of 5 μL and incubated on ice for one hour before separation by native gel electrophoresis. SDS-free sample buffer was added, and 1 % acrylamide gels equilibrated in chilled 0.5 x TBE buffer were used to separate the indicated reaction products. DNA was visualized with SYBR Gold Nucleic Acid Gel Stain (ThermoFisher) according to the manufacturer’s instructions. Nucleosome concentration was estimated using absorbance at 260 nM. Titrations were carried out to determine the minimal concentration that could be visualized using SYBR Gold stain, and this was ∼100 nM for all experiments shown. Mif2 kinase treatments were carried out as described for inner kinetochore reconstitution experiments, and total NaCl concentration was normalized to 108 mM for every binding reaction.

### In vitro *kinetochore assembly assay*

Plasmid pRS316 and pRS316-*CEN6Δ* (HZE3047) were used to generate *CEN6* (348 bp) and control DNA (586 bp) templates, respectively. To make biotinylated DNA templates, PCR reactions were performed by using the following primers:

Biotinylated 5’ primer: 5’-bio-CAGGAAGGCAAAATGCCGC

Unlabeled 3’ primer (*CEN6*): 5’-GTATTTGTTGGCGATCCCCC

Unlabeled 3’ primer (control DNA): 5’-GATCGCTTGCCTGTAACTTA

Twelve 50 μL PCR products were pooled, precipitated in ethanol, gel-purified, and conjugated to Streptadivin-coated Dynabeads (M-280 Streptavidin, Invitrogen) for 1 hour at room temperature. To ensure the beads have equivalent numbers of DNA substrates (∼14 pmol/ 100 μL beads), 3 μg of *CEN6* DNA and 5 μg of control DNA were conjugated to 100 μL of streptavidin beads, respectively. Beads were washed twice and stored in Buffer L (25 mM Hepes-K, pH 8.0; 175 mM K-glutamate; 15% Glycerol; 2 mM MgCl_2_; 0.1 mM EDTA; 0.5 mM EGTA, 0.1% NP-40) at 4°C until use.

Kinetochore assembly was done according to a previous report with minor modifications (Lang *et al*., 2018). Briefly, cells expressing Flag-Cse4 with Mif2-TAF, Mif2-10A-TAF, or Mif2-10D-TAF were grown in 1 liter of YPD to an OD_600_ of 1.2 and harvested by centrifugation. Cells were washed once with buffer L, and then the cell pellets were resuspended in buffer L supplemented with protease inhibitors (2 mM PMSF; 200 μM benzamidine; 0.5 μg/ml leupeptin; and 1 μg/ml pepstatin A) before being snap-frozen in liquid nitrogen drop-by-drop. Cells were then lysed by cryogenic milling using a Freezer/Mill (SPEX SamplePrep). Cell powder was thawed on ice and clarified as described above. The concentration of clarified whole cell extracts (WCE) was between 50-60 mg/mL, which were used freshly or stored at -80°C for later use.

For each kinetochore assembly assay, 150 μL of WCE was incubated with 10 μL of DNA-loaded beads on a rotator at room temperature for 45 min. Beads were washed three times with 1 mL cold buffer L before being boiled in 20 μL of 1x LDS sample buffer (Invitrogen) supplemented with 50 mM DTT. WCE and bound proteins were separated by running in a 4-12% Bis-Tris gradient gel with MOPS SDS running buffer (Invitrogen). Mif2-TAF and Flag-Cse4 were detected by immunoblotting with anti-protein A (Sigma, P3775, 1:10,000) and anti-Flag (Sigma, F3165, 1:2000) antibodies, respectively.

### ChIP-qPCR assay

To evaluate the localization of Mif2, Mif2-10A, and Mif2-10D to the centromere, ChIP was performed (Meluh and Koshland, 1997; Suhandynata *et al*., 2019). Briefly, yeast cultures (150 mL, enough for three immunoprecipitation experiments) were grown to an OD_600_ of 0.8 and cross-linked for 15 min with 1 % formaldehyde at room temperature. Whole-cell lysates were prepared in 0.8 ml ChIP lysis buffer (50 mM Hepes, pH 7.6; 140 mM NaCl; 1 mM EDTA; 1 % Triton, and 0.1% sodium deoxycholate) supplemented with protein inhibitors by glass bead beating and sonication to shear the genomic DNA to an average size of 300–500 bp. Immunoprecipitation was performed using 50 μL Dynabeads (Protein G; ThermoFisher) and 3 μL anti-Flag antibody M2 (Sigma; F3165). After binding, beads were washed as follows: once in 1 mL lysis buffer with 5 min incubation, twice in 1 mL washing buffer (100 mM Tris-Cl, pH 8.0; 250 mM LiCl; 0.5% NP-40; 0.5% deoxycholate; and 1 mM EDTA) with 5 min incubations, and once with 1 mL TE buffer (10 mM Tris-Cl, pH 8.0, and 1 mM EDTA) with 1 min incubation. After the washes, the samples were first eluted in 40 μL of TE buffer with 1% SDS at 65°C for 10 min, and this was saved as elution 1. 162 μL of DNA extraction buffer (135 μL of TE, 15 μL of 10% SDS, 12 μL of 5 M NaCl) was then added to the beads with 1.5 μL of RNaseA and incubated at 37°C for 30min, and this was elution 2. Both eluates were mixed and incubated at 37 °C for another 30 min. The inputs were treated with 162 μL of DNA extraction buffer and 1.5 μL of RNaseA with 1 h incubation at 37°C.

The input and immunoprecipitated DNA were incubated in the same buffer with the addition of 20 μg Protease K at 65 °C overnight to reverse crosslinks before purification using a QIAquick PCR Purification kit (QIAGEN). Before qPCR analysis, the input DNA was diluted 1:100, and immuno-purified DNA was diluted 1:10 by volume. qPCR was done using SYBR Green 2x master mix (KAPA Biosystems) on a Roche LightCycler 480 system. Three independent immunoprecipitation experiments were performed. qPCR primer sequences were:

CEN3_fwd: 5′-ATCAGCGCCAAACAATATGGAAAA-3′

CEN3_rev: 5′-GAGCAAAACTTCCACCAGTAAACG-3′

CUP1_fwd: 5′-AACTTCCAAAATGAAGGTCA-3′

CUP1_rev: 5′-GCATGACTTCTTGGTTTCTT-3′.

### Co-immunoprecipitation and Mif2 competition experiments

To detect the *in vivo* association of endogenous Mif2-WT/10A with the Cse4 nucleosome, we used strains expressing Mif2-TAF variants and tagged Cse4 (internal 3xFLAG) from their respective chromosomal loci. The 3xFLAG tag was inserted at the XbaI site of Cse4, similar to a previous report (Wisniewski et al., 2014). Mif2-TAF/FLAG-Cse4 or *mif2-10A*-TAF/FLAG-Cse4 cells were harvested by centrifugation and washed (PBSN, 1 mM EDTA, 1 mM NaF, 1 mM beta-glycerophosphate, PI and PMSF). For each IP, 80 mL of OD_600_ 1 cells were resuspended in 4x cell pellet volumes of lysis buffer (∼0.8 mL PBSN, 0.2% NP-40) and frozen dropwise in liquid nitrogen before lysis using a freezer mill. The resulting powder was thawed at 4 °C before clarification by centrifugation, and the protein concentration of each cell extract was normalized to ∼10 mg/mL. 0.8 mL of clarified lysate was incubated with 30 μL of IgG beads (human IgG, ∼3 mg Protein A per mL capacity) at 4°C overnight with rotation. The beads were then washed with 1mL of ice-cold lysis buffer four times before elution of bound material via a 5 min incubation at 98 °C with 30 μL of 2x LDS buffer (Invitrogen). About 25 μL of eluate was collected by centrifugation. 2 μL of 1 M DTT was added to each sample before heating at 98 °C for 2 min to reduce the protein samples.

To evaluate the stability of the pre-existing Mif2-Cse4 binding against competition by newly synthesized Mif2-WT, Mif2-10A and Mif2-10D proteins, we transformed the Mif2-TAF/FLAG-Cse4 strain with high-copy 2-micron pRS425 plasmids coding for galactose-inducible Mif2-3xHA, Mif2-10A-3xHA, or Mif2-10D-3xHA. The Mif2-3xHA variants were induced by addition of 2% galactose to log phase cultures grown in SC-leu/raf (SC-leu with 2 % raffinose instead of glucose). For these experiments, cells at three time points were collected: pre-induction, one hour after induction, and two hours after induction. The co-IP method described above was used to evaluate endogenous Mif2-Cse4 binding in the presence of Mif2 competitors.

Input and eluted samples were loaded onto a NuPAGE gradient gel (4-12 %; ThermoFisher) and run in MOPS-SDS buffer (ThermoFisher) at 200 V for 70 min. After transferring at 100V for 100 min to a PVDF membrane, the membrane was blotted with anti-Protein A (1:10,000, Sigma), anti-HA (1:2000, 3F10, Sigma), and anti-FLAG (1:2000, Sigma) primary antibodies to detect Mif2-TAF, Mif2-3xHA, and FLAG-Cse4, respectively.

---

**Supplementary file 1.**
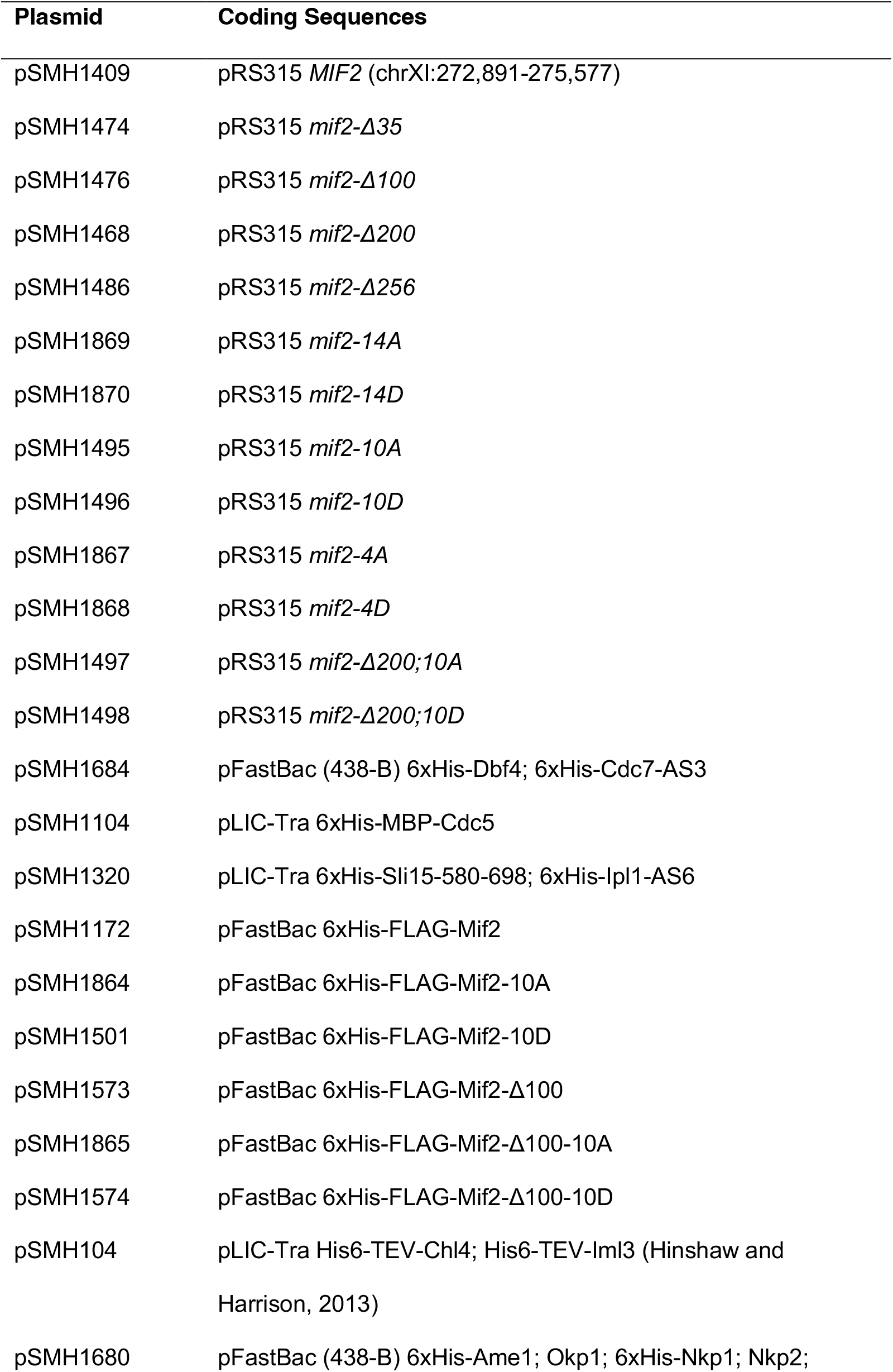
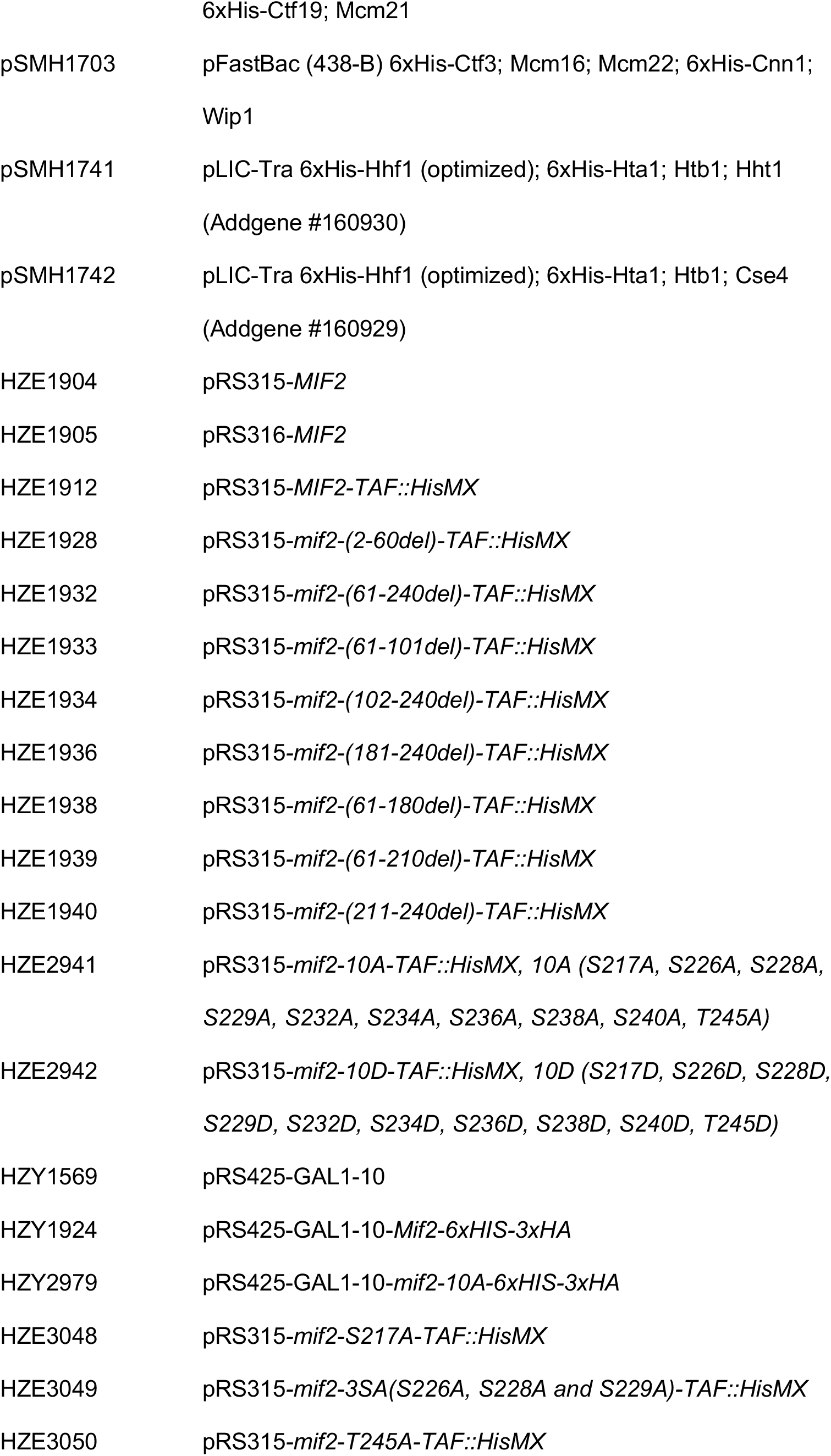
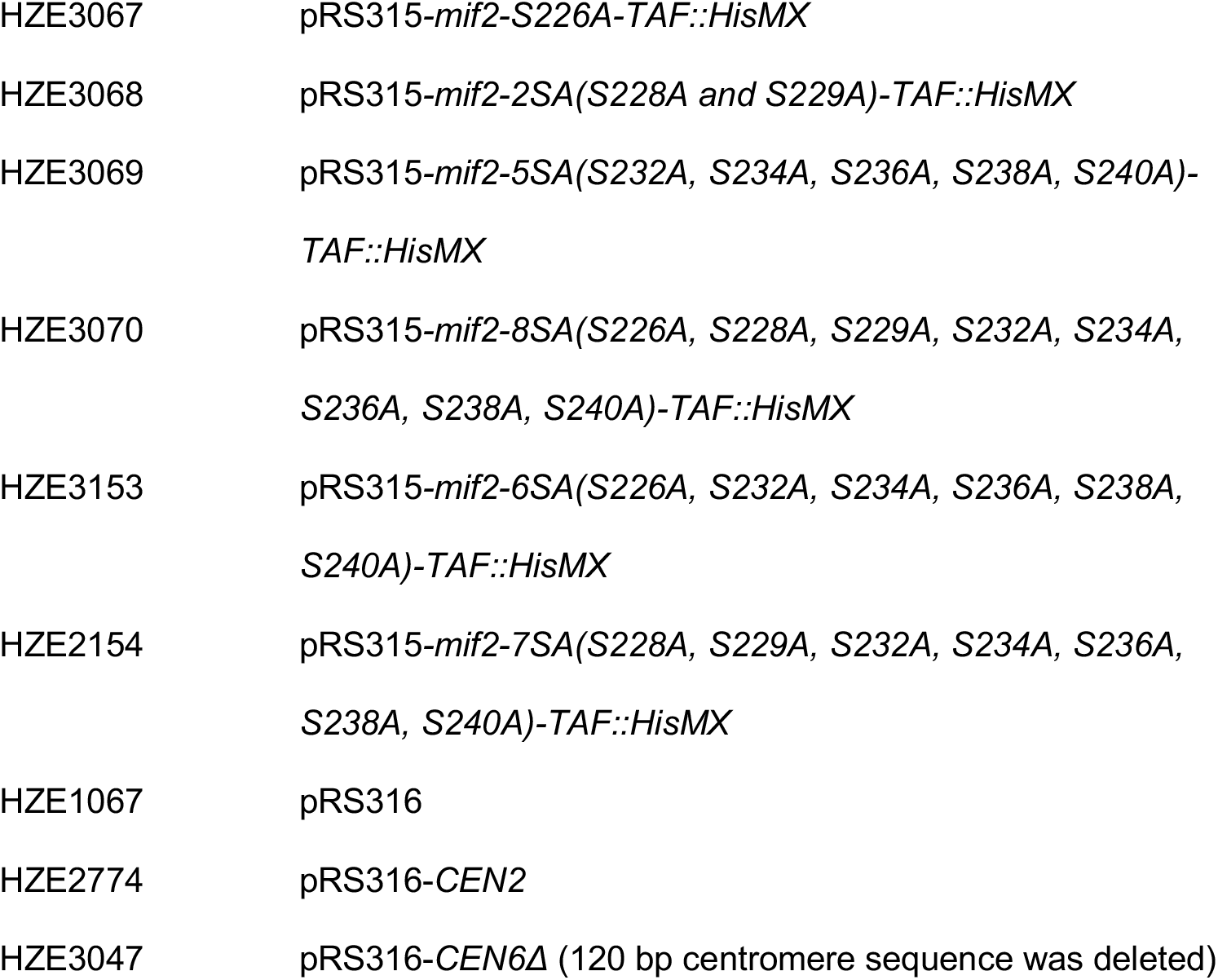
Plasmids used in this work

**Supplementary file 2.**
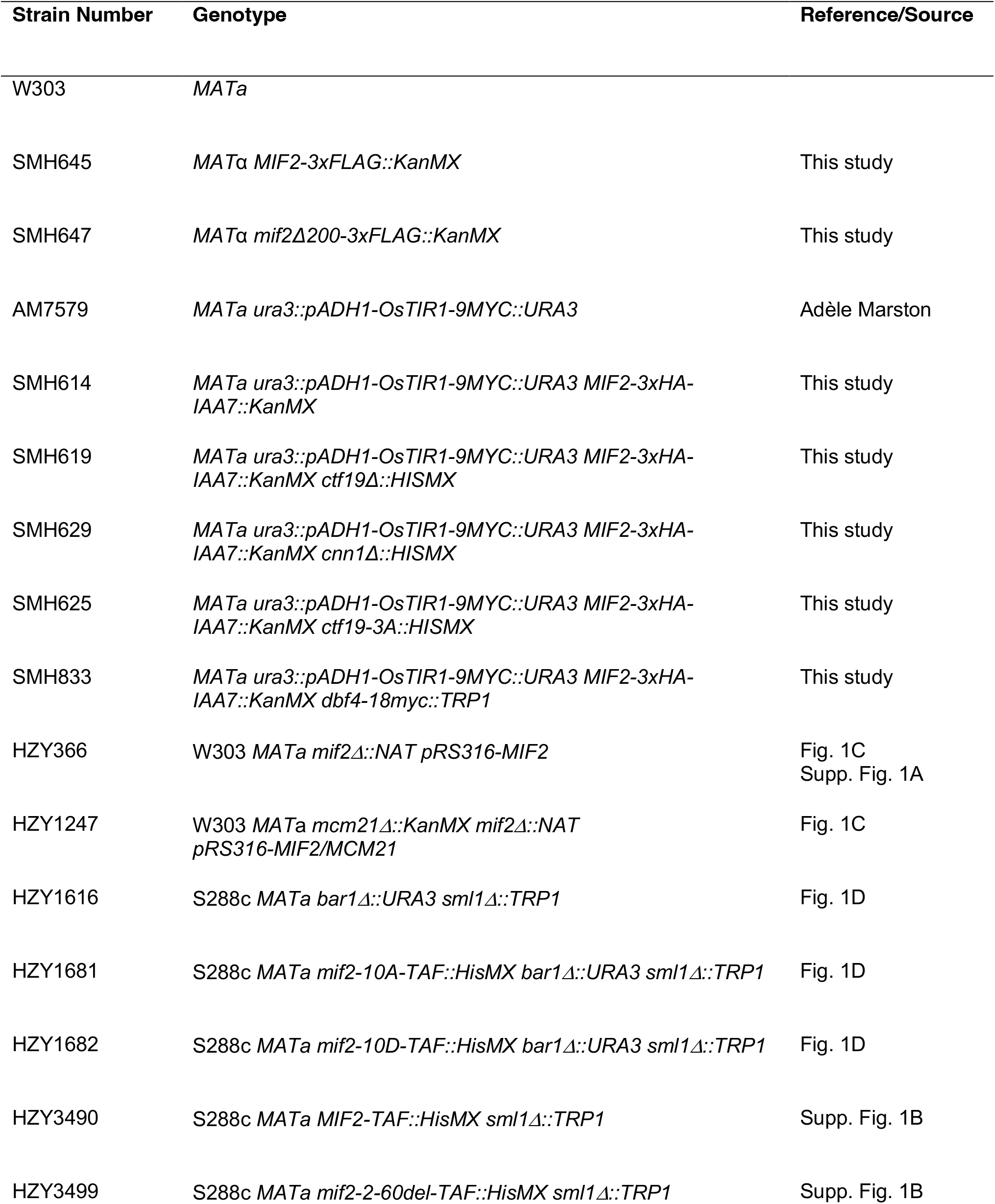
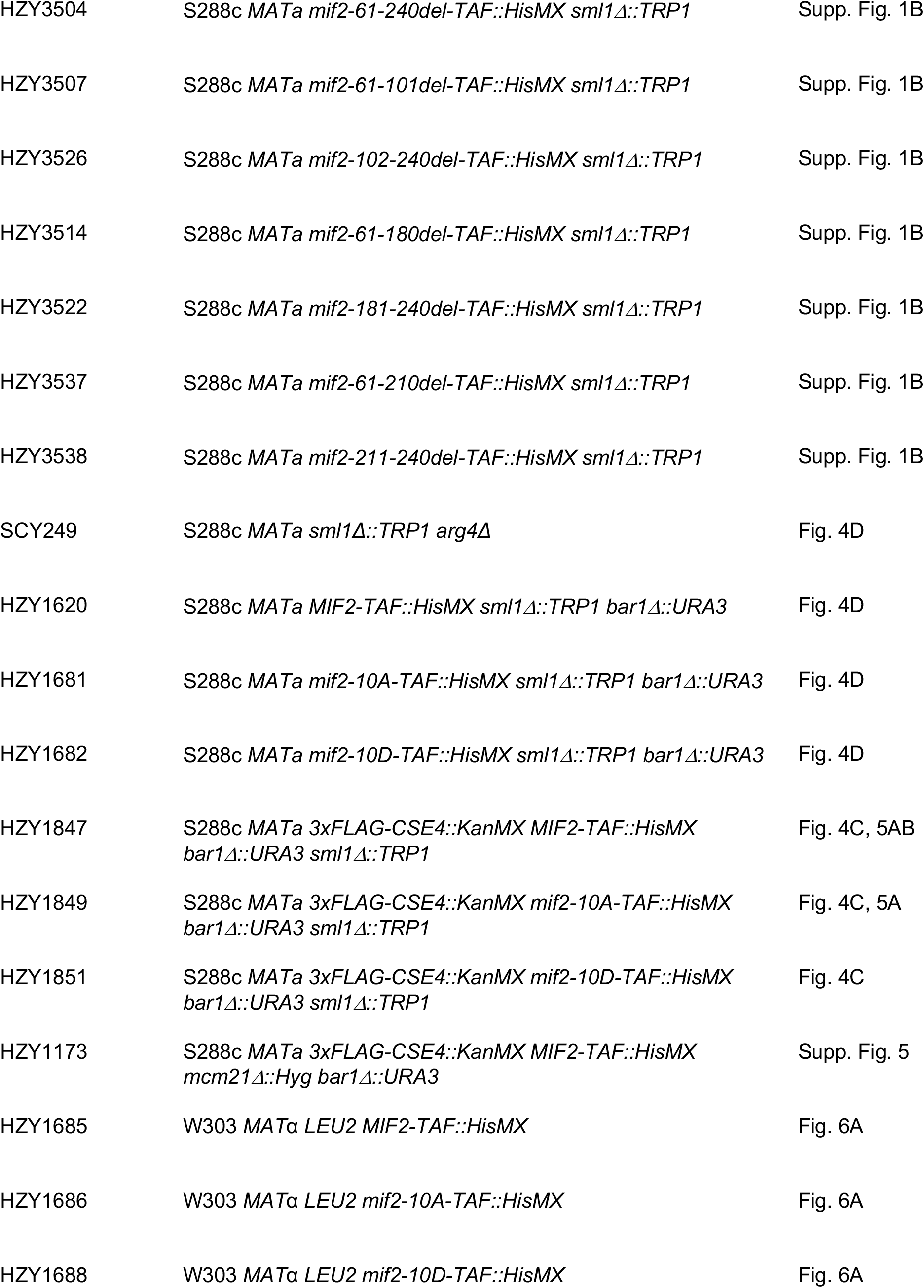
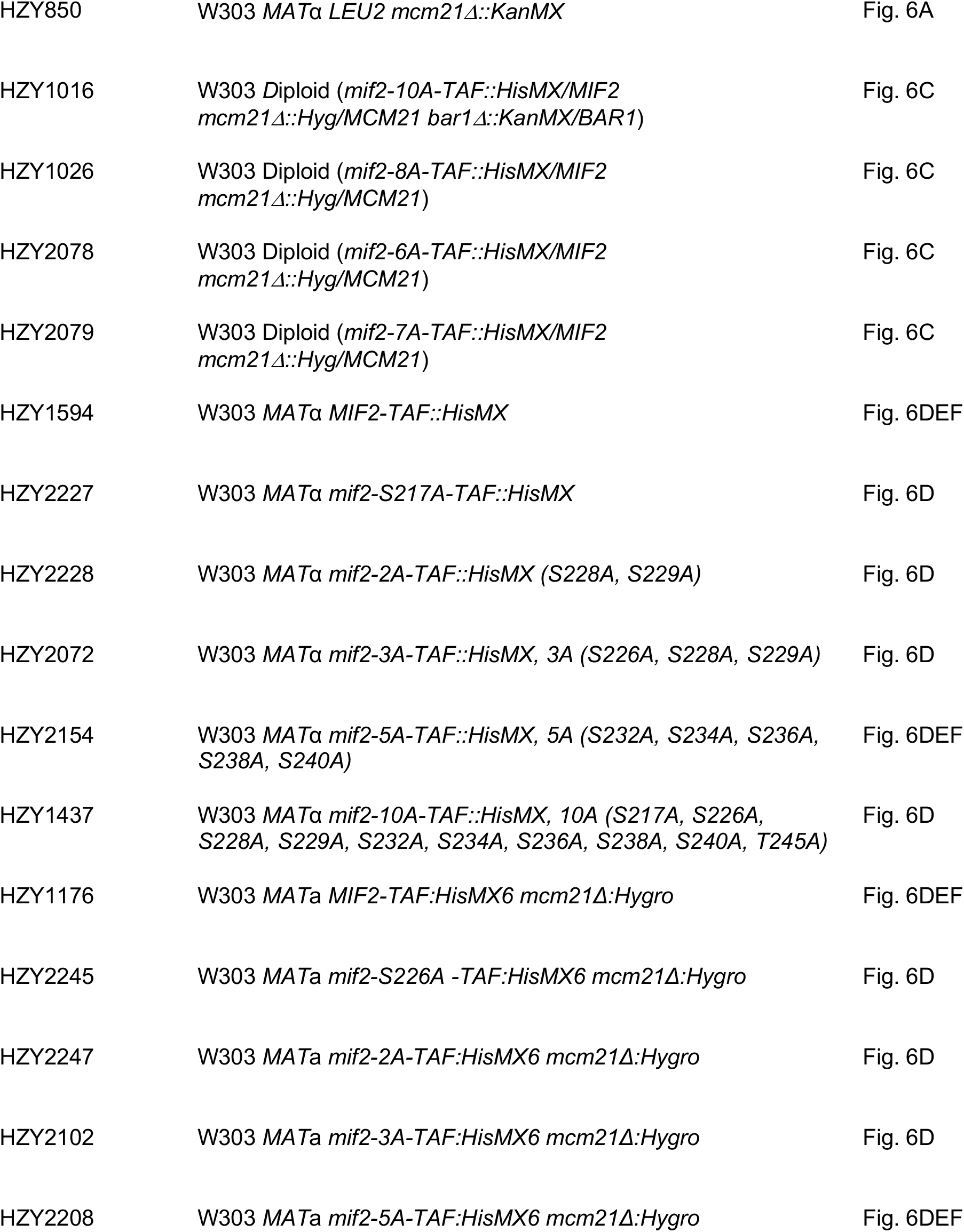
Yeast strains used in this study

